# Combinative protein expression of immediate early genes c-Fos, Arc, and Npas4 along aversive- and reward-related neural networks

**DOI:** 10.1101/2025.04.21.649441

**Authors:** Mary Arai, Hisayuki Osanai, Chris C. Snell, Takashi Kitamura, Sachie K. Ogawa

## Abstract

Expression of immediate early genes (IEGs) is critical for memory formation and has been widely used to identify the neural substrate of memory traces, termed memory engram cells. Functions of IEGs have been known to be different depending on their types. However, there is limited knowledge about the extent to which different types of IEGs are selectively or concurrently involved in the formation of memory engram. To address this question, we investigated the combinative expression of c-Fos, Arc, and Npas4 proteins using immunohistochemistry following aversive and rewarding experiences across subregions in the prefrontal cortex (PFC), basolateral amygdala (BLA), hippocampal dentate gyrus (DG), and retrosplenial cortex (RSC). Using an automated cell detection algorithm, we found that expression patterns of c-Fos, Npas4, and Arc varied across different brain areas, with a higher increase of IEG expressing cells in the PFC and posterior BLA than in the DG. The combinative expression patterns, along with their learning-induced changes, also differed across brain areas; the co-expression of IEGs increased in the PFC and BLA following learning whereas the increase was less pronounced in the DG and RSC. Furthermore, we demonstrate that different area-to-area functional connectivity networks were extracted by different IEGs. These findings provide insights into how different IEGs and their combinations identify engram cells, which will contribute to a deeper understanding of the functional significance of IEG-tagged memory engram cells.

## 1. Introduction

Formation of long-term memory requires transcription and protein synthesis (Asok et al 2019, Barondes & Jarvik 1964, Dash et al 1990, Davis & Squire 1984, Flexner et al 1963, Kandel 2001, Silva et al 1998). Immediate-early genes (IEGs) have rapid and transient transcription upon extracellular stimulation (Greenberg & Ziff 1984, Gu et al 2023, Sheng & Greenberg 1990). IEGs are involved in long-term synaptic plasticity and memory, and are widely used as endogenous markers of neuronal activity, animal’s experience, and pharmacological activation (Barth et al 2004, Fuentes-Ramos & Barco 2024, Guzowski et al 1999, Hoffman et al 1993, Minatohara et al 2015, Okuno 2011, Salery et al 2021, Yokose et al 2024, Yokose et al 2023). Furthermore, IEGs have been utilized for the identification of memory engram cells, which are subpopulations of neurons that are activated by a salient experience and subsequently undergo biological changes to encode a specific memory episode (Josselyn et al 2015, Josselyn & Tonegawa 2020, Kandel et al 2014, Liu et al 2012a, Silva et al 2009). Transgenic approaches using IEGs have enabled the identification of memory engram cells through visualizing and optogenetic/chemogenetic manipulation of IEG-expressing cells allowing tracking the memory-bearing cells and investigating their causal roles in learning and memory (Barth et al 2004, Choi et al 2018, Denny et al 2014, Kitamura et al 2017, Liu et al 2012a, Marks et al 2022, Ortega-de San Luis & Ryan 2022, Reijmers et al 2007, Tanaka et al 2018, Terranova et al 2022, Terranova et al 2023, Tonegawa et al 2015, Vetere et al 2019, Wang et al 2006, Yamamoto et al 2021). The expression of IEGs has been considered to be involved in synaptic plasticity by neuronal activity (Tonegawa et al 2015), and importantly, different types of IEG have different roles in synaptic plasticity and memory. For example, c-Fos, a transcription factor IEG (Morgan et al 1987, Sagar et al 1988, Yap & Greenberg 2018), is essential for synaptic plasticity and memory consolidation (Fleischmann et al 2003, Katche et al 2010, Kemp et al 2013), with increased dendritic spine density in expressing cells (Choi et al 2018, Ryan et al 2015, but see Uytiepo et al 2025). Arc (activity-regulated cytoskeletal protein), an effector IEG (Guzowski et al 1999, Nikolaienko et al 2018), influences long-term memory (Plath et al 2006) by heterosynaptically weakening inactive synapses (El-Boustani et al 2018, Minatohara et al 2015, Okuno et al 2012, Yap & Greenberg 2018). Npas4 (neuronal PAS domain protein 4), another transcription factor IEG (Lin et al 2008), is crucial for memory consolidation (Ramamoorthi et al 2011, Weng et al 2018) and regulates excitatory-inhibitory synaptic balance (Spiegel et al 2014, Sun & Lin 2016). However, difference in the type of IEG has been relatively not considered in identifying memory engram cells, because there is still limited knowledge about the extent to which different IEGs are selectively or concurrently involved in engram cell formation.

Several studies have reported the co-expression (Gonzales et al 2020, Stone et al 2011) and segregation (Sun et al 2020, Ye et al 2016) of different IEGs after learning. However, those investigations are limited to specific IEG, behavior, and brain region. The aim of this study is to address the extent to which different types of IEGs are selectively or concurrently expressed in individual cells in multiple brain regions simultaneously after different experiences. Using the automated cell detection method we previously proposed (Osanai et al 2025), we investigated the combinative protein expression of c-Fos, Arc, and Npas4 with IHC after aversive and reward experience across the mPFC, basolateral amygdala (BLA), hippocampal dentate gyrus (DG), and retrosplenial cortex (RSC), which are implicated in aversive and appetitive memory (Giustino & Maren 2015, Gore et al 2015, Gourley & Taylor 2016, Kesner 2018, Kheirbek et al 2013, Kirk et al 2017, Kitamura et al 2017, Kwapis et al 2015, Redondo et al 2014, Sierra-Mercado et al 2011, Sun et al 2021, Terranova et al 2022, Vedder et al 2017).

## 2. Materials and methods Animals

All procedures relating to mouse care and experimental treatments conformed to NIH and Institutional guidelines, and were conducted with the approval of the UT Southwestern Institutional Animal Care and Use Committee (IACUC). Total 18 male C57BL/6J mice between 8–16 weeks old were used. Mice were group housed with littermates (2–5 mice per cage) in a 12-hour light/dark cycle until a day before experiments.

### Sample preparation and Imaging

We prepared brain section samples with three different conditions: home cage (HC), Contextual fear conditioning (CFC), and Reward conditioning (RC) groups (n = 6 for each group, three to six sections per animal). Mice had ad libitum access to food and water except the RC group.

#### Home cage

Mice were separated into individual cages 1 day before the sampling, and then deeply anesthetized with a ketamine (75 mg/kg)/dexmedetomidine (1 mg/kg) (K/D) cocktail for transcardial perfusion with 4% paraformaldehyde (PFA) in PBS. Brains were removed and post-fixed in 4% PFA in PBS at 4°C at least for 24 hours.

#### Contextual fear conditioning

Mice were separated into individual cages 1 day before CFC. Foot-shock stimulation was provided based on the previous reports (Osanai et al 2025, Osanai et al 2023, Terranova et al 2022). In this study, a fear apparatus with a 24 cm W × 20 cm D × 20 cm H chamber (Med Associates) was used, and a mouse was placed in the fear stimulation chamber for a 3-minute habituation period and for subsequent 3-minute shock period. During the shock period, the mouse received three foot shocks (0.75 mA, 2-sec) with 58-second inter-shock intervals. After the stimulation, the mouse was returned to its home cage for one hour. The mouse was then immediately anesthetized deeply with a K/D cocktail and transcardially perfused with 4% PFA in PBS. Brains were removed and post-fixed in 4% PFA in PBS at 4°C at least for 24 hours. The shock chamber was cleaned before starting each experiment.

#### Reward conditioning

Mice were separated into individual cages and subjected to one week food-restriction with access to a small amount of food daily, resulting in a reduction of their body weight to ∼85% of the initial weight. The RC protocol was conducted in the same chamber as the CFC. After the food restriction period, each mouse was allowed to explore the chamber with a food pellet for 30 minutes. Eating behavior during chamber exploration was confirmed through video recording and by measuring the weight reduction of the provided food pallet. The mouse was then returned to its home cage for one hour. Following this, the mouse was immediately anesthetized deeply with a K/D cocktail and transcardially perfused with 4% PFA. Brains were removed and post-fixed in 4% PFA in PBS at 4°C at least for 24 hours. The chamber was cleaned before starting each experiment.

#### Immunohistochemistry and Imaging

The fixed brains were sectioned using a vibratome (Leica VT100S) with a thickness of 60 μm. For immunohistochemistry (IHC), tissue sections were washed with PBS, blocked with 0.03% Triton-X PBS (PBS-T) with 5% normal donkey serum (NDS) (Jackson ImmunoResearch Labs; RRID: AB_2337258) for 30 minutes, and then incubated with primary antibodies diluted in the PBS-T with 5% NDS for two nights at 4°C. After washing with PBS (3×5 min), tissue sections were subsequently incubated with secondary antibodies in the PBS-T with 5% NDS for 2 hours at room temperature. Primary antibodies were chicken anti-NeuN (1/1000, Millipore Sigma, ABN91; RRID: AB_11205760), guinea pig anti-Arc (1/500, synaptic systems, 156005; RRID: AB_2151848), rabbit anti-Npas4 (1/1000, Activity signaling, AS-AB18A-300), and goat anti-cFos (1/1000, Santacruz, sc-52-G; RRID:AB_2629503). Secondary antibodies were donkey anti-chicken DyLight 405 (1/500, Jackson ImmunoResearch Labs; RRID: AB_2340373), donkey anti-guinea pig AlexaFluor488 (1/500, Jackson ImmunoResearch Labs; RRID: AB_2340472), donkey anti-rabbit AlexaFluor546 (1/500, ThermoFisher Scientific; RRID: AB_2534016), and donkey anti-goat AlexaFluor633 (1/500, Jackson ImmunoResearch Labs; RRID: AB_2535739). After incubating in the secondary antibody solution, the tissue sections were washed in PBS (2×5 min) and mounted in VECTASHIELD antifade mounting medium (Vector Laboratories) on glass slides.

All fluorescence images (0.624 µm/pixel) were acquired under the same imaging condition using Zeiss LSM800, 10× objective lens (NA: 0.45), and Zen Blue software (Zeiss). Details of the imaging conditions were described previously (Osanai et al 2025).

### Analysis

IEG- and NeuN-positive cells were detected in the prelimbic (PL) and infralimbic (IL) regions of PFC, anterior and posterior BLA (aBLA and pBLA), granule cell layers of dorsal and ventral DG (dDG and vDG), and dorsal/ventral parts of anterior and posterior RSC (dorsal aRSC, ventral aRSC, dorsal pRSC, and ventral pRSC), whose boundaries were determined based on the Allen Brain Reference Atlas (Allen Institute for Brain Science 2004). From bregma, PL and IL were determined in the coronal sections at +1.845 to +1.42 mm, aBLA was at -1.255 to -1.655 mm, pBLA was at -2.355 to -2.78 mm, dDG was at -1.655 to -2.255 mm, vDG was at -3.28 to -3.455 mm, aRSC was at -1.255 mm to -2.255 mm, and pRSC was at -2.78 mm to -3.78 mm. The regions of interest (ROI) for each brain region were manually drawn using ImageJ (Schindelin et al 2012). The images of each channel (NeuN, c-Fos, Npas4, Arc) and the ROI information were imported into MATLAB R2024b (Mathworks) for further analysis. For the analysis in the PL and dDG, we included the data used in the previous report (Osanai et al 2025).

#### Automated cell detection

In this study, we used the newly developed method, automated cell detection after background assumption (ADABA) algorithm, written in MATLAB that we proposed previously (Osanai et al 2025). Briefly, the algorithm subtracted the background of the image and then the background-cleaned image was used for cell detection. First, the images were converted into 8-bit gray-scale and median filters were applied (c-Fos, Npas4, and Arc images: 11x11 pixels; NeuN image: 7x7 pixels). For assuming background pattern, we first determined the intense signal pixels in the image by drawing 20 intensity contours using Otsu’s method (Otsu 1979) which is implemented in MATLAB Image Processing Toolbox (*multithresh.m*). A contour which covers more than 80% (for Arc and NeuN images) or 95% (for c-Fos and Npas4 images) of the image was selected for further calculation. The inside-contour areas were filled with the neighboring intensity, and the background pattern was assumed by filtering with spatial moving average filter (31x31 pixels). The assumed background was then subtracted from the median-filtered image. Then, thresholding was conducted on the background-subtracted image with T = k * σ, where T is the threshold, σ is standard deviation intensity of the background-subtracted image, and k is coefficient parameter; k = 5 for c-Fos, Npas4, and Arc, and k = 2 for NeuN positive cell detections. The thresholded signals were denoised and smoothed by morphological operations of erosion-reconstruction with five pixels distance and closing with two pixels distance. Signals whose areas are smaller than 50 pixels were regarded as noise and removed for further processing. The smoothed thresholded signals were then segmented with a watershed algorithm to detect c-Fos, Npas4, Arc, and NeuN positive cells in the image. IEGs were assessed as co-localized within a cell if their detected areas share more than five pixels. Peak fluorescent intensity of each detected cell was measured to evaluate the expression level of IEGs.

Similar to the previous report (Osanai et al 2025), to evaluate the accuracy of the automated cell detection, manual cell detection was performed by an experimenter that was blinded to the result of the automated cell detection using ImageJ (Schindelin et al 2012). For manual detection, random 77 images of HC, 129 images of CFC, and 128 images of RC of the PL, IL, aBLA, pBLA, dDG, and vDG were used. The Precision, or false positive detection rate, was calculated as Precision = TP / (TP + FP), where true positive (TP) and false positive (FP) were manually checked after the automated detection. To evaluate the ratio that cells detected manually were also detected in the automated algorithm, the Auto-Manual match rate, or Sensitivity, was calculated as Match / (Match + Eye_Only), where Match indicates the number of cells detected both by the automated and manual approach and Eye_Only indicates the number of cells identified only by manually. To evaluate the ratio of cells overlooked in manual identification but detected in the automated algorithm, the Sensitivity-increase rate was calculated as Auto_Only / (Match + Auto_Only) where Auto_Only indicates the number of cells that were not identified manually but detected in the automated algorithm. The custom code used in this study is available at https://github.com/HisayukiOsanai/CellDetection.

#### Network analysis

To calculate IEG-based functional connectivity networks (Silva et al 2019, Takeuchi et al 2022, Tanimizu et al 2017, Vetere et al 2017, Wheeler et al 2013), correlation matrices of Pearson r values were calculated between all 10 brain regions within each experimental group (HC, CFC, RC) and each IEG-group (c-Fos, Npas4, Arc, and combinations of them). Hierarchical clustering of the correlation matrix was visualized by average-linkage hierarchical clustering with dissimilarity index, or distance, calculated by distance = 1 - |r| (Liu et al 2012b, Takeuchi et al 2022).

Functional connectivity networks were constructed by thresholding the correlation matrices and visualized using MATLAB. The network connection lines, or edges, represent Pearson correlations |r| between brain areas. A correlation of |r| > 0.7 was considered strong and used for thresholding the network connections. Line thickness and node sizes are proportional to the |r| value between brain areas and to the number of connections each brain area has, individually. Complexity of networks was evaluated by Connectivity per brain area which indicates the number of connections (edges) per brain area (node), and by Connectivity per effective node which indicates the number of connections (edges) per brain area that has at least one connection (effective node). To evaluate dissimilarity between graphs, graph-edit-distance (GED) (Bai et al 2019, Tantardini et al 2019, Wills & Meyer 2020) and Sum of Differences in Edge-Weight Values (SDEWV) (Wang et al 2019) were calculated. GED is a measurement of the minimum number of operations required to transform one graph into another, which we calculated by the sum of all elements of the difference between two graph adjacency matrices: GED = ∑_i,j_ D_i,j_ , where D_i,j_ is the element of difference matrix **D** = **A** – **A’**, with **A** and **A’** being the adjacency matrices of the two graph **G** and **G’**. SDEWV is the sum of absolute differences in edge weights between two graphs, which we calculated as SDEWV = ∑_i,j_ |e_i,j_ - e^1^ |, where e and e’ are edge weight between nodes *i* and *j* in the two graphs.

#### Statistics

Statistical analyses were performed using MATLAB. All bar plots and error bars represent mean ± standard error, and box plots in the violin plots display minimum, 25th percentile, median, 75th percentile, and maximum values. One-way ANOVA followed by post-hoc Tukey test was used to analyze differences between groups. For cell-intensity analysis, we used nonparametric Kruskal-Wallis test followed by Dunn-Šidák test. Effect sizes were calculated using Cohen’s d, where d = 0.2, 0.5, 0.8, and < 0.2 are considered as small, medium, large, and negligible effects (Cohen 1988, Thomas et al 1991). For effect sizes in the cell-intensity analysis, we calculated Cliff’s δ, where δ = 0.147, 0.33, 0.474, and < 0.147 correspond to small, medium, large, and negligible effects (Cliff 1993, Macbeth et al 2011, Meissel & Yao 2024, Romano et al 2006). p < 0.05 was considered to be statistically significant. *, **, and *** indicate p < 0.05, 0.01, and 0.001, respectively. Due to the large sample size in the cell-intensity analysis, we assessed non-negligible difference using effect size rather than relying solely on p-values in the intensity analysis (Nakagawa & Cuthill 2007). #, ##, and ### indicate δ ≥ 0.147, 0.33, and 0.474, respectively, with all p < 0.05. Unless otherwise noted, n = 6 animals for each group.

## 3. Results

### Automated detection of c-Fos, Npas4, and Arc expressing cells

For the comprehensive investigation of IEG-expressing cells in multiple brain regions, we used the automated cell detection method. Brain slice images were captured using the confocal microscope following the IHC staining with c-Fos-, Npas4-, and Arc-antibodies. (Fig. 1A). Consistent with our previous results (Osanai et al 2025), the automated cell detection has high accuracy compared to manual detection; the false positive detection was very low (Precision > 0.96), and the automated detection identified more than 77% of the manually identified cells with increased sensitivity in the confocal images of the BLA, PFC and DG (Fig. 1B). Some IEG-stained images contained both strongly and weakly labeled cells, making consistent manual cell counting difficult; cells with similar intensity were more likely to be counted manually in images with sparse cell distribution but were often overlooked in images with dense and variable-intensity staining cells (Fig. 1A yellow rectangles) (Osanai et al 2025). Such observation bias can be avoided by automated detection, resulting in higher sensitivity of Arc positive cell detection (Fig. 1B right). The number of detected cells was highly correlated with R > 0.7 between automated and manual cell detections for all c-Fos, Npas4, and Arc positive cells (Fig. 1C). The results indicate that our automated cell detection algorithm has high precision and helps reduce observation bias in the manual detection of IEG expressing cells.

**Figure 1:**
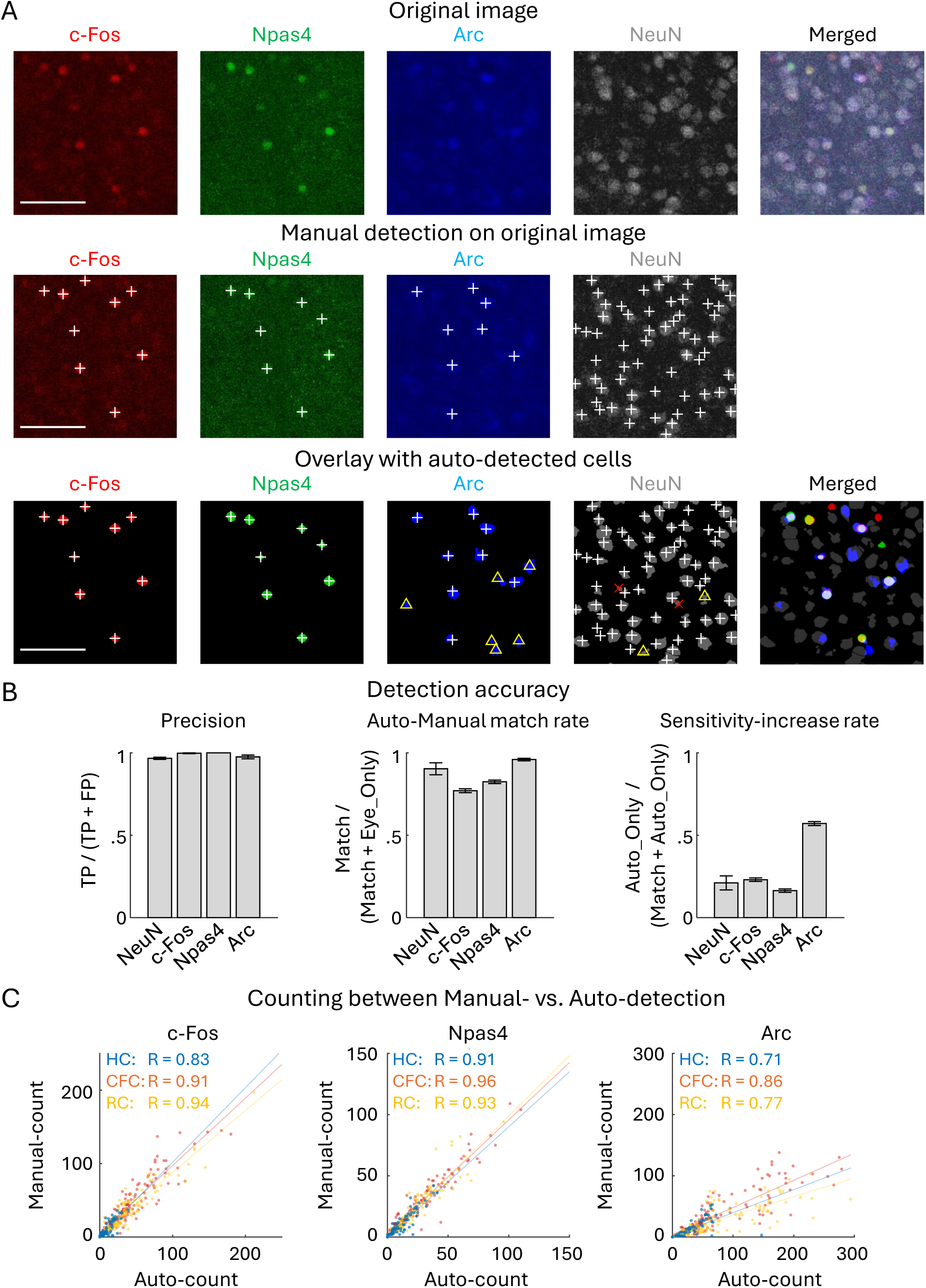
Automated detection of IEG positive cells and the detection accuracy (A) Top: Original images of c-Fos, Npas4, Arc, and NeuN expressed cells in aBLA. Middle: Overlay of the manually detected positive cells (white ‘+’ marks) on the original images. Bottom: Auto-detected cells (colored cells), overlay of the manually detected positive cells and manually verified false-positive detections (red ‘×’ marks). Yellow triangles indicate cells overlooked in the manual detection but detected in the automated algorithm. Scale bars, 100 µm. (B) Evaluation of auto-detection accuracy with Precision, Auto-Manual match rate, and Sensitivity increase rate. Precision indicates false positive ratio in the automated cell detection per total detected cells, that is Precision = TP / (TP + FP), where true positive (TP) and false positive (FP) were identified manually after the automated detection (n = 7, 9, 7, 9 sections for NeuN, c-Fos, Npas4, and Arc). Auto-Manual match rate indicates the ratio of cells which were both manually and automatically detected, such that Auto-Manual match rate = Match / (Match + Eye_Only), where Match indicates the number of cells which were both manually and automatically detected, and Eye_Only indicates the number of cells identified only by manual detection (n = 5 sections for NeuN, n = 334 sections for c-Fos, Npas4, and Arc). Sensitivity increase rate indicates ratio that cells were not detected manually but detected automatically, that Sensitivity-increase rate = Auto_Only / (Match + Auto_Only), where Auto_Only indicates the number of cells which were not identified manually but detected automatically (n = 5 sections for NeuN, n = 334 sections for c-Fos, Npas4, and Arc). (C) Correlation of automatically and manually detected cell number, with R indicating Pearson correlation coefficient (p < 0.001 for all). HC: n = 77, CFC: n = 129, RC: n = 128 sections. The sections used for accuracy analysis were randomly selected from the PFC, BLA, and DG.

### c-Fos, Npas4, and Arc expression in various brain areas

Due to the reliability of automated cell detection, we applied this method to identify the positive cells for all following analyses. The cell detections for c-Fos, Npas4, and Arc were performed in the prelimbic cortex (PL) (Fig. 2, S1A), infralimbic cortex (IL) (Fig. 2, S1B), anterior BLA (aBLA) (Fig. 2, S2A), posterior BLA (pBLA) (Fig. 2, S2B), dorsal DG (dDG) (Fig. 2, S3), ventral DG (vDG) (Fig. 2, S4), dorsal/ventral anterior RSC (aRSC) (Fig. 2, S5), and dorsal/ventral posterior RSC (pRSC) (Fig. 2, S6), individually 60 min following CFC or reward conditioning (RC) (see Methods). After CFC, c-Fos^+^ cell density was significantly increased in all brain areas except the ventral aRSC compared to the home-cage (HC) group. On the other hand, Npas4^+^ cell density was increased in the PL, IL and pBLA. The Arc^+^ cell density was increased in the PL, IL, pBLA, vDG, ventral aRSC, and dorsal/ventral pRSC (Fig. 3A, S7, S8, S9A). After RC, the c-Fos^+^ cell densities were increased significantly in all brain areas; Npas4^+^ cell densities were increased in the PL, IL, pBLA, and dorsal/ventral aRSC; Arc^+^ cell densities were increased in the PL, IL, pBLA, dDG, dorsal aRSC, and dorsal/ventral pRSC (Fig. 3A, S7, S8, S9A). Furthermore, after CFC, the average expression level of c-Fos was increased in the PL, IL, a/pBLA, d/vDG, and ventral pRSC; Npas4 expression level was increased in the aBLA and decreased in the dorsal aRSC and dorsal/ventral pRSC; Arc expression level increased in the PL, aBLA, and dDG and decreased in the ventral pRSC. After RC, the average c-Fos expression level was increased in all brain areas but the ventral aRSC; Npas4 expression level was increased in the PL and pBLA; Arc expression level increased in the aBLA, dDG, ventral aRSC, and ventral pRSC (Fig. 3B, S7, S8, S9B). The decrease of average expression level after stimulation in RSC may indicate that the number of cells with weak IEG expression were increased whereas the maximum expression level is strictly controlled compared to other brain areas once IEG is expressed. Thus, the expressions of c-Fos, Npas4, and Arc are differently increased after the conditioning experience depending on the brain areas (Fig. 3A). However, overall, brain areas with a greater increase in the expression of one IEG type tended to show higher expression of other types of IEGs with correlation coefficient larger than 0.7 (Fig. S10).

**Figure 2:**
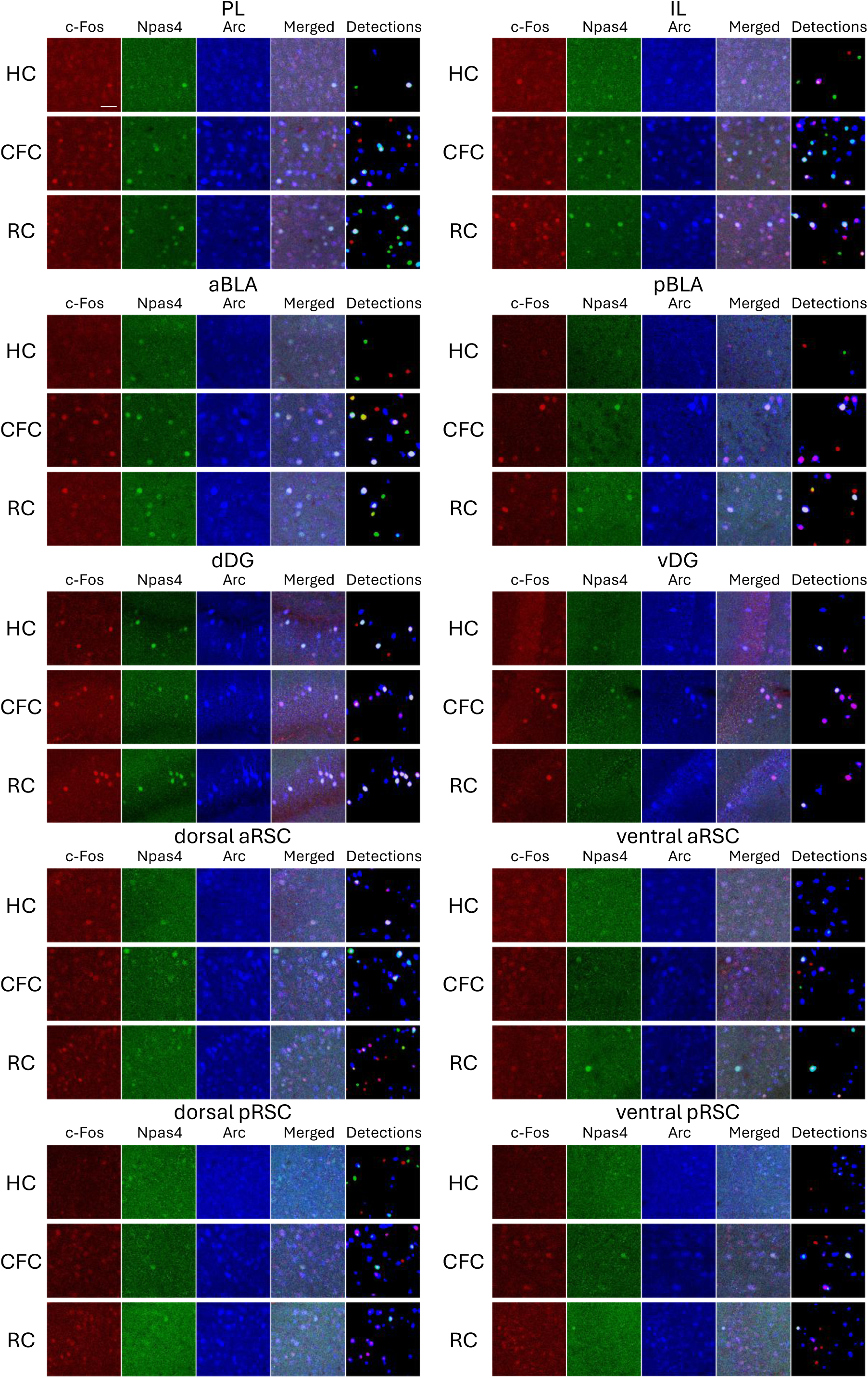
Expression of c-Fos, Npas4, and Arc in multiple brain regions. Images of c-Fos (red), Npas4 (green), Arc (blue), merged, and auto-detected cells in the prelimbic (PL), infralimbic (IL), anterior basolateral amygdala (aBLA), posterior BLA (pBLA), dorsal dentate gyrus (dDG), and ventral DG (vDG), dorsal anterior RSC (aRSC), ventral aRSC, dorsal posterior RSC (pRSC), and ventral pRSC. Top: Home cage (HC); Middle: Contextual fear conditioning (CFC); Bottom: Reward conditioning (RC) groups. Scale bars, 40 µm. Lager field-of-view images are shown in Supp. Figure S1–S8.

**Figure 3:**
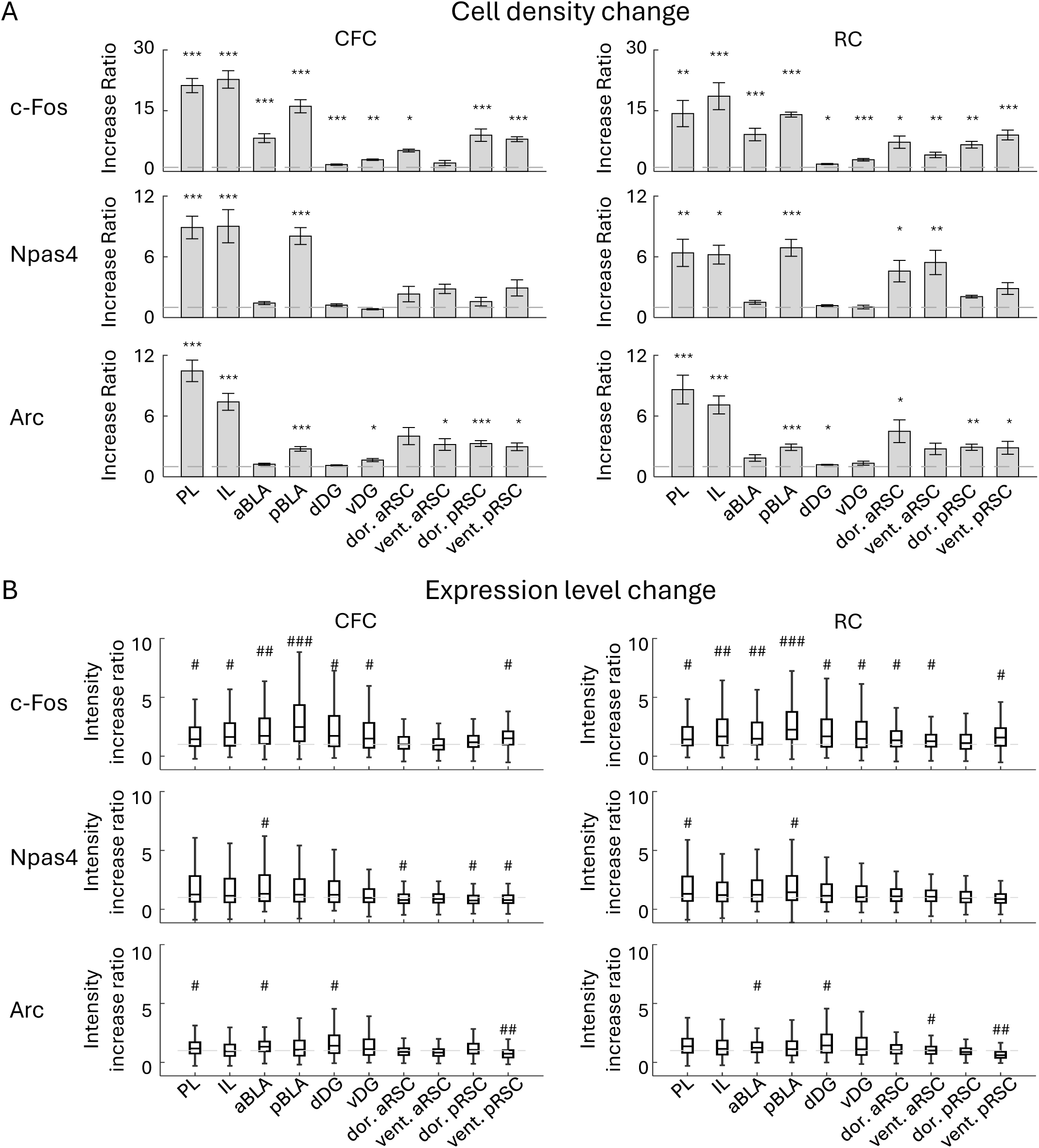
Changes of cell densities and expression levels of c-Fos, Npas4, and Arc positive cells. (A) Fold changes of c-Fos, Npas4, and Arc positive cell densities by CFC and RC in each brain region, compared with HC group. (B) Fold changes of c-Fos, Npas4, and Arc expression levels of cells by CFC and RC in each brain region, compared with HC group. Statistical tests were conducted between groups, as shown in Supp. Figure S7 and S8.

### Combinative expression of cFos, Npas4, and Arc

Next, we investigated the extent to which the different types of IEGs are selectively- or co-expressed in individual neurons. In both PL and IL, cell densities of the double-positive cells of different IEGs, and c-Fos/Npas4/Arc triple-positive cells tended to be increased by mouse experience of conditioning (Fig. 4A, E). The ratio of IEG single-positive cells was decreased in both PL and IL by CFC or RC, but instead the ratio of the c-Fos/Npas4/Arc triple-positive cells and double-positive cells of c-Fos/Arc and Npas4/Arc were increased (Fig. 4B, C, F, G; S11A, B). The expression levels of c-Fos, Npas4, and Arc in single cells in both PL and IL were not correlated in the HC group, but the correlation between c-Fos and Npas4 and between Npas4 and Arc increased after CFC and RC (Fig. 4D, H). These indicate that c-Fos, Npas4, and Arc tend to be independently expressed in the HC condition, but become co-expressed after CFC and RC in the PL and IL.

**Figure 4:**
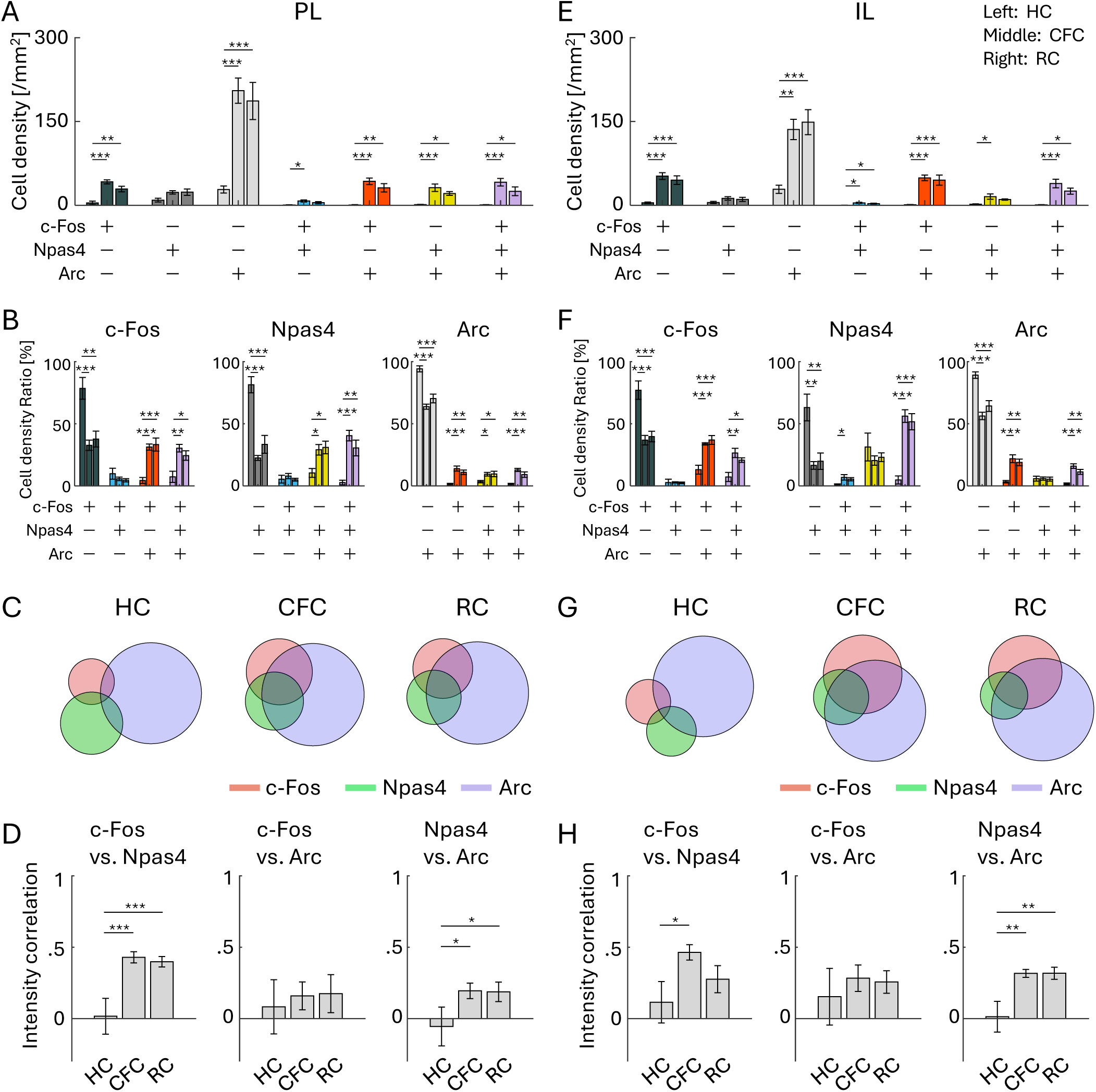
Co-expression of IEGs in PL and IL. (A, E) Cell densities of each cell group with selective or combinative IEG expression in the PL (A) and IL (E). (B, F) Ratio of cell densities per all c-Fos cells (left), Npas4 cells (middle), and Arc cells (right), in the PL (B) and IL (F). (C, G) Venn diagrams of c-Fos, Npas4, and Arc positive cells in HC, CFC, and RC groups, in the PL (C) and IL (G). The size of the circles corresponds to cell densities, normalized by Arc cell density. (D, H) Average correlation of IEG expression in single cells, between c-Fos vs. Npas4 (left), c-Fos vs. Arc (middle), and Npas4 vs. Arc (right), in the PL (D) and IL (H). For each group of bars, the left bar indicates HC, the middle indicates CFC, and the right indicates RC.

In the aBLA, cell densities of the c-Fos/Npas4/Arc triple-positive cells and c-Fos/Arc double-positive cells were increased by CFC and RC (Fig. 5A). Also, the ratio of the c-Fos/Npas4/Arc triple-positive cells was increased by CFC and RC (Fig. 5B, C; S11C). The expression levels of IEGs were correlated after CFC and RC but not in HC (Fig. 5D). Similarly, in the pBLA, cell density of the c-Fos/Npas4/Arc triple-positive cells was increased by CFC and RC, as well as c-Fos/Npas4 and c-Fos/Arc double-positive cells and c-Fos single-positive cells (Fig. 5E). The ratio of the c-Fos/Npas4/Arc triple-positive cells was increased by CFC and RC (Fig. 5F, G; S11D). The correlation of expression level between c-Fos and Npas4 was significantly increased after CFC and RC compared to HC (Fig. 5H). These results indicate that, similar to the results in the PFC, c-Fos, Npas4, and Arc tend to be co-localized after CFC and RC in the BLA.

**Figure 5:**
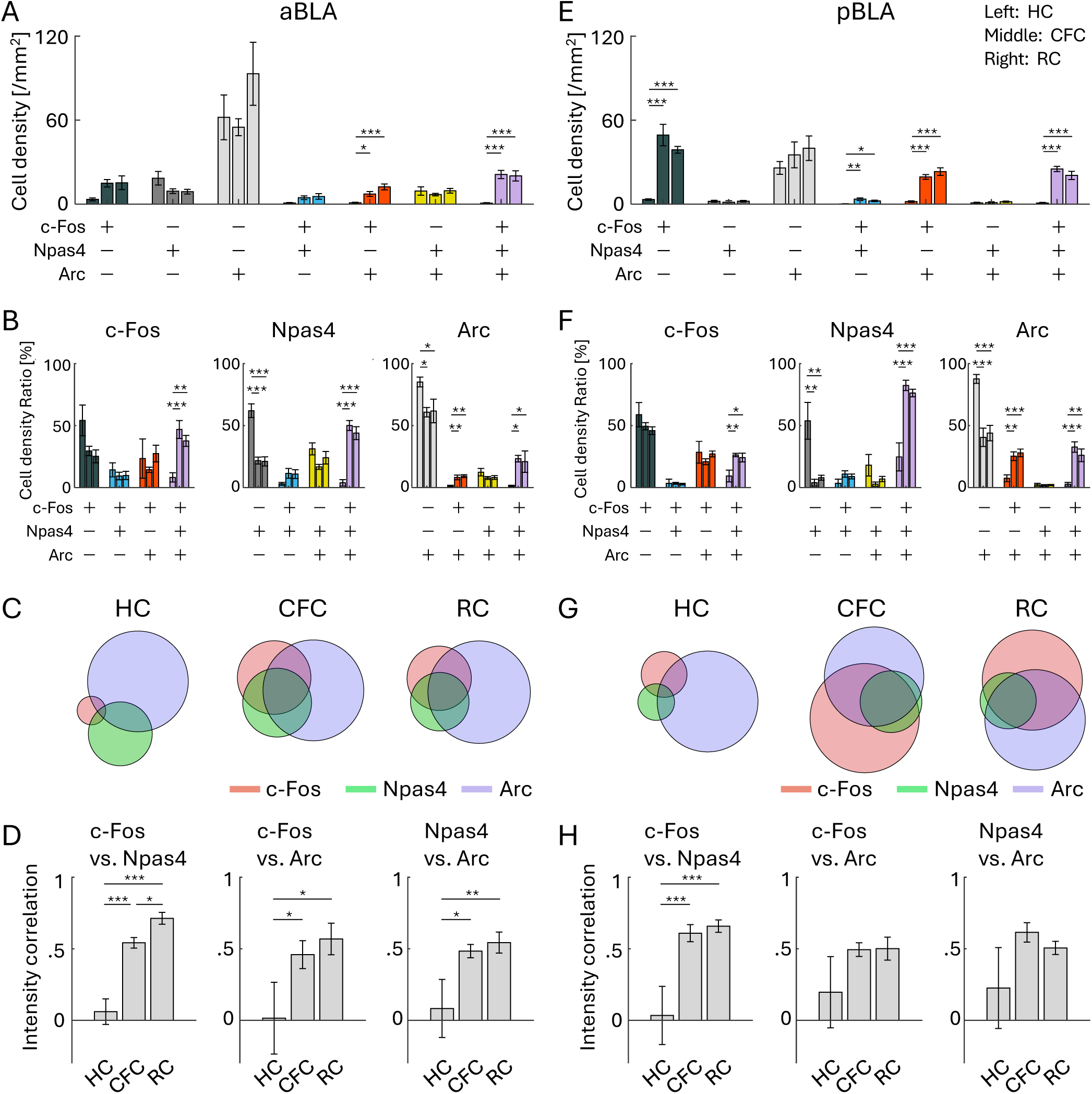
Co-expression of IEGs in anterior and posterior BLA. (A, E) Cell densities of each cell group with selective or combinative IEG expression in aBLA (A) and pBLA (E). (B, F) Ratio of cell densities per all c-Fos cells (left), Npas4 cells (middle), and Arc cells (right), in the aBLA (B) and pBLA (F). (C, G) Venn diagrams of c-Fos, Npas4, and Arc positive cells in HC, CFC, and RC groups, in the aBLA (C) and pBLA (G). The size of the circles corresponds to cell densities, normalized by Arc cell density. (D, H) Average correlation of IEG expression in single cells, between c-Fos vs. Npas4 (left), c-Fos vs. Arc (middle), and Npas4 vs. Arc (right), in the aBLA (D) and pBLA (H). For each group of bars, the left bar indicates HC, the middle indicates CFC, and the right indicates RC.

On the other hand, in the DG, cell density of the c-Fos/Npas4/Arc triple-positive cells was increased by CFC and RC in the dDG but the change was not observed in the vDG (Fig. 6A, E). Unlike the PFC and BLA, where the proportion of the triple-positive cells was <3% of whole population in the HC group (Fig. S11A-D), both dDG and vDG had a higher ratio of triple-positive cells in HC (26.7 ± 3.0% in dDG and 12.3 ± 1.7% in vDG; Fig. S11E, F) as well as CFC and RC groups (Fig. 6B, C, F, G; S11E, F). The increase of the triple-positive cell ratio after CFC and RC was only observed within Npas4^+^ cells but not within c-Fos^+^ and Arc^+^ cells in the dDG (Fig. 6B), and the increase was not observed in the vDG (Fig. 6F, G). Unlike the PFC and BLA, CFC and RC groups did not show increased correlations of expression level in any pair between c-Fos, Npas4, and Arc in the dDG and vDG compared to HC (Fig. 6D, H). Thus, the combinative expression patterns of c-Fos, Npas4, and Arc in the dDG and vDG were different from the PFC and BLA.

**Figure 6:**
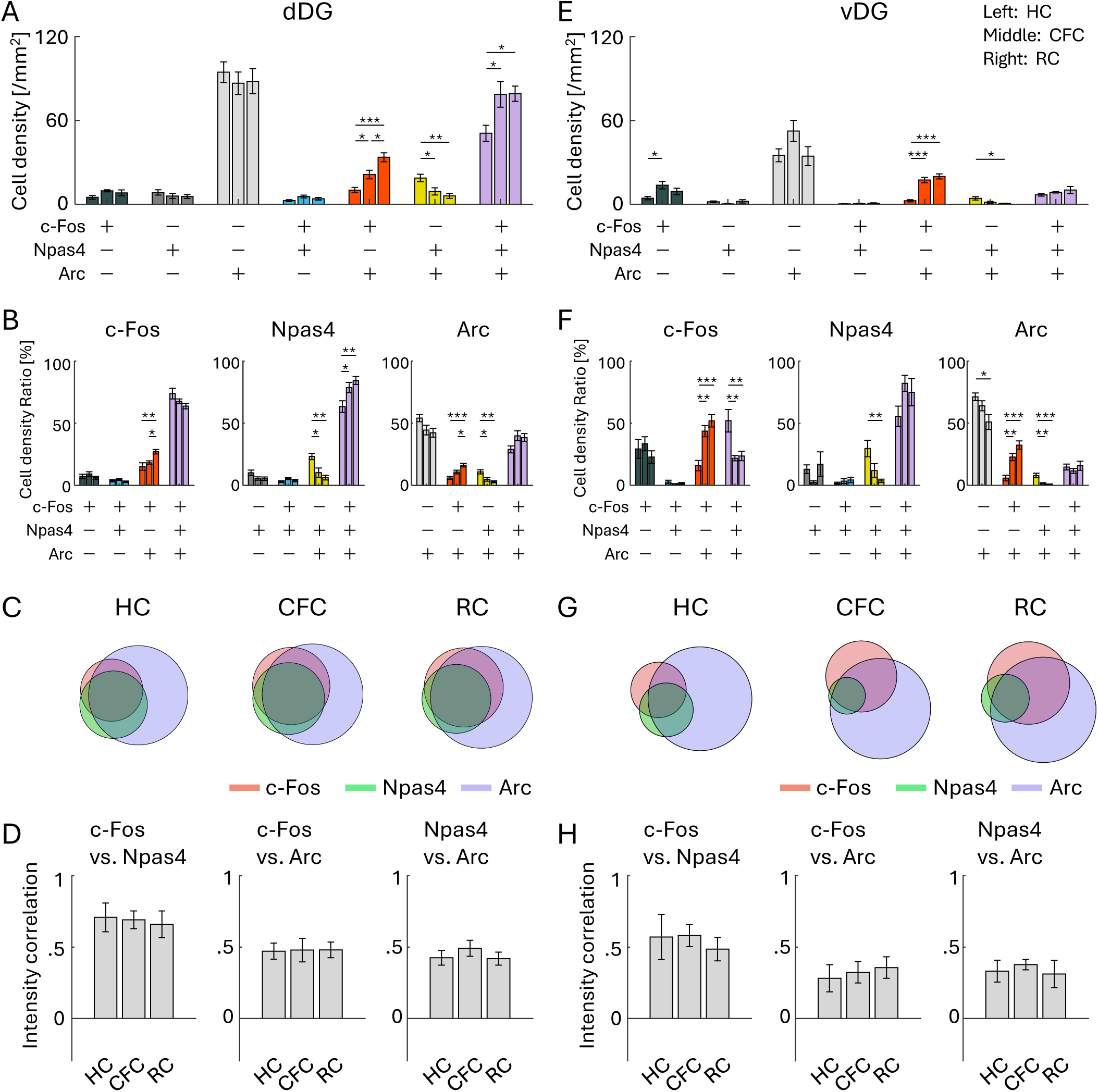
Co-expression of IEGs in dorsal and ventral DG. (A, E) Cell densities of each cell group with selective or combinative IEG expression in the dDG (A) and vDG (E). (B, F) Ratio of cell densities per all c-Fos cells (left), Npas4 cells (middle), and Arc cells (right), in the dDG (B) and vDG (F). (C, G) Venn diagrams of c-Fos, Npas4, and Arc positive cells in HC, CFC, and RC groups, in the dDG (C) and vDG (G). The size of the circles corresponds to cell densities, normalized by Arc cell density. (D, H) Average correlation of IEG expression in single cells, between c-Fos vs. Npas4 (left), c-Fos vs. Arc (middle), and Npas4 vs. Arc (right), in the dDG (D) and vDG (H). For each group of bars, the left bar indicates HC, the middle indicates CFC, and the right indicates RC.

Lastly, we investigated IEG colocalization in the aRSC and pRSC. In the aRSC, cell density of the c-Fos/Npas4/Arc triple-positive cells was significantly increased by RC but not clearly observed by CFC both in the dorsal and ventral aRSC (Fig. 7A, E). The increase in the ratio of the triple-positive cells was not clear in the dorsal aRSC (Fig. 7B, C; S12G), and only significant within Npas4^+^ and Arc^+^ cells in the ventral aRSC (Fig. 7F, G; S11H). Increase of the correlation of c-Fos, Npas4, and Arc expression levels was not clear in dorsal/ventral aRSC (Fig. 7D, H). In the pRSC, cell density of the c-Fos/Npas4/Arc triple-positive cells was significantly increased by CFC and RC in the dorsal pRSC (Fig. 7I), and by CFC in the ventral pRSC (Fig. 7M). The increase in the ratio of the triple-positive cells was not significant in dorsal pRSC (Fig. 7J, K; S11I), and was significant within c-Fos^+^ and Npas4^+^ cells in the ventral pRSC (Fig. 7N, O; S11J). Increase of the correlation of c-Fos, Npas4, and Arc expression levels was not clear in dorsal/ventral pRSC (Fig. 7L, P). Therefore, the increase of the c-Fos/Npas4/Arc triple-positive cells in the aRSC/pRSC was less clear than the PFC and BLA, but was distinct from the DG.

**Figure 7:**
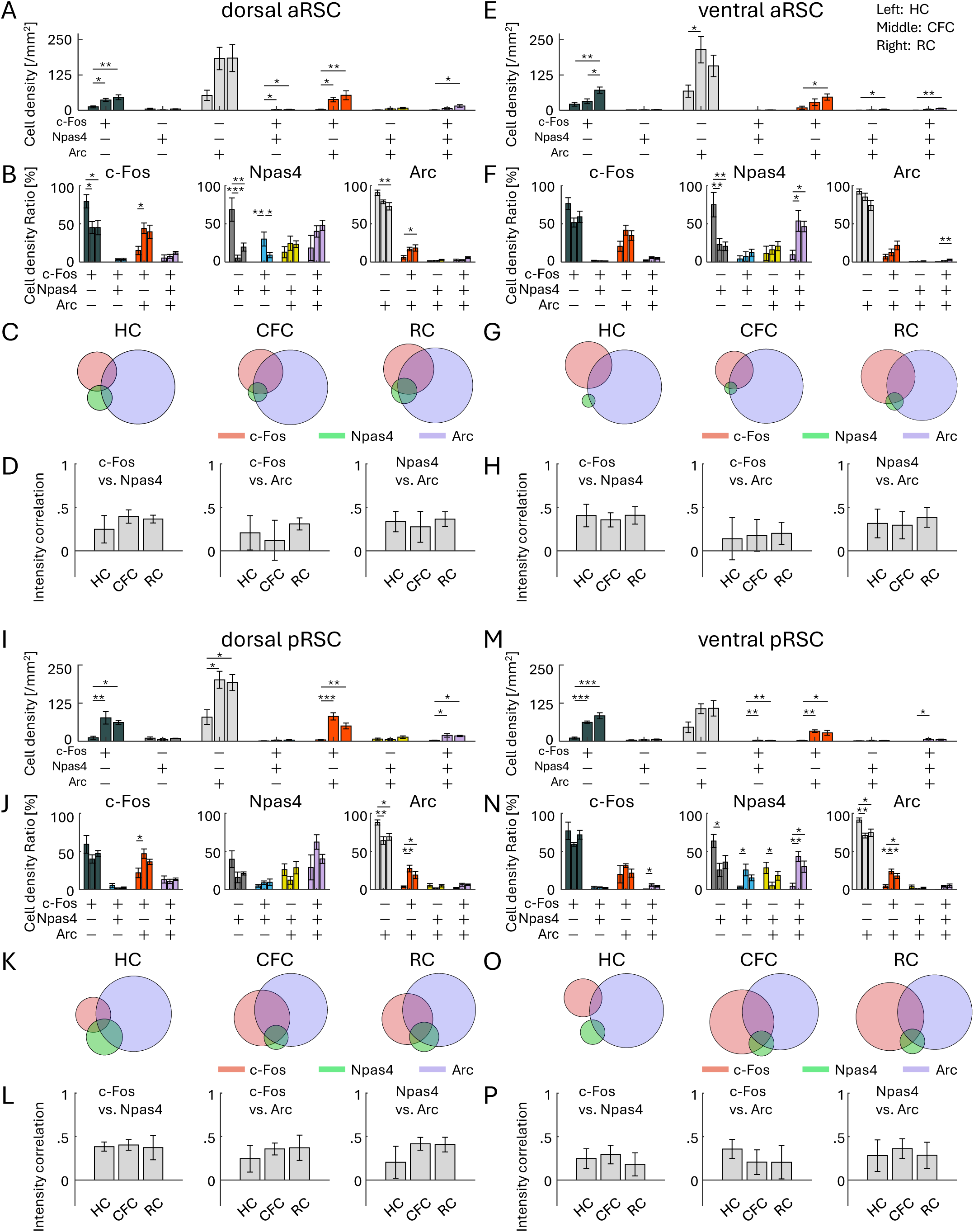
Co-expression of IEGs in RSC. (A, E, I, M) Cell densities of each cell group with selective or combinative IEG expression in the dorsal aRSC (A), ventral aRSC (E), dorsal pRSC (I), and ventral pRSC (M). (B, F, J, N) Ratio of cell densities per all c-Fos cells (left), Npas4 cells (middle), and Arc cells (right), in the dorsal aRSC (B), ventral aRSC (F), dorsal pRSC (J), and ventral pRSC (N). (C, G, K, O) Venn diagrams of c-Fos, Npas4, and Arc positive cells in HC, CFC, and RC groups, in the dorsal aRSC (C), ventral aRSC (G), dorsal pRSC (K), and ventral pRSC (O). The size of the circles corresponds to cell densities, normalized by Arc cell density. (D, H, L, P) Average correlation of IEG expression in single cells, between c-Fos vs. Npas4 (left), c-Fos vs. Arc (middle), and Npas4 vs. Arc (right), in the dorsal aRSC (D), ventral aRSC (H), dorsal pRSC (L), and ventral pRSC (P). For each group of bars, the left bar indicates HC, the middle indicates CFC, and the right indicates RC.

We have also investigated expression levels of c-Fos, Npas4, and Arc in each c-Fos/Npas4/Arc positive/negative cell group (Fig. S12, S13, S14). The increases or decreases population expression levels were observed, but we could not find systematic tendencies (Fig. S15, S16, S17) . Also, we could not find systematic tendencies in correlation changes of IEG expression levels in different cell groups (Fig. S18, S19, S20).

Altogether, we observed brain area-specific changes of combinative expression of cFos, Npas4, and Arc induced by CFC or RC (Table 1, 2). The degree of combinative IEGs expression increase varied across areas. Interestingly, the DG did not show a clear increase of the c-Fos/Npas4/Arc triple-positive ratio after CFC and RC in contrast to the PFC, BLA, and RSC.

**Table 1.**
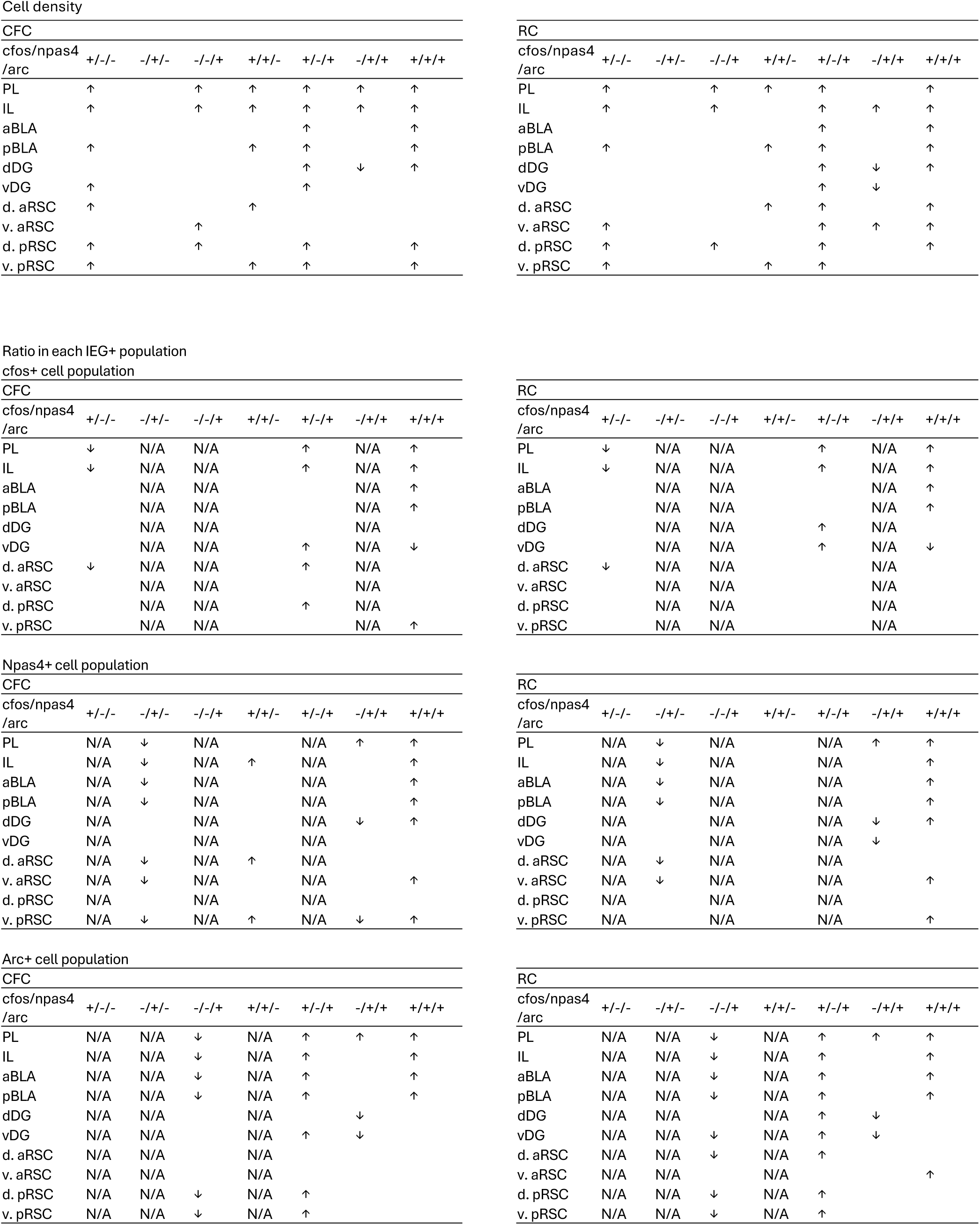
Changes in IEG overlapping cells. Summary of changes in cell densities of each cell group and cell density ratio per each IEG type, induced by CFC and RC. Arrows indicate p < 0.05. Cohen’s d were >0.8 for all.

**Table 2.** Changes in expression level correlation between IEGs. Summary of changes in expression level correlation between c-Fos, Npas4, and Arc in individual cells, induced by CFC and RC. Arrows indicate p < 0.05. Cohen’s d were >0.8 for all.

| CFC |  |  |  |
| --- | --- | --- | --- |
|  | c-Fos vs.<br>Npas4 | c-Fos vs.<br>Arc | Npas4 vs.<br>Arc |
| PL | ↑ |  | ↑ |
| IL | ↑ |  | ↑ |
| aBLA | ↑ | ↑ | ↑ |
| pBLA | ↑ |  |  |
| dDG |  |  |  |
| vDG |  |  |  |
| d. aRSC |  |  |  |
| v. aRSC |  |  |  |
| d. pRSC |  |  |  |
| v. pRSC |  |  |  |

| RC |  |  |  |
| --- | --- | --- | --- |
|  | c-Fos vs.<br>Npas4 | c-Fos vs.<br>Arc | Npas4 vs.<br>Arc |
| PL | ↑ |  | ↑ |
| IL |  |  | ↑ |
| aBLA | ↑ | ↑ | ↑ |
| pBLA | ↑ |  |  |
| dDG |  |  |  |
| vDG |  |  |  |
| d. aRSC |  |  |  |
| v. aRSC |  |  |  |
| d. pRSC |  |  |  |
| v. pRSC |  |  |  |

### IEG expression-based area-area connectivity

Since IEGs have also been employed to examine the functional connectivity between brain areas based on the correlation of IEG expression (Franceschini et al 2023, Silva et al 2019, Takeuchi et al 2022, Tanimizu et al 2017, Vetere et al 2017, Wheeler et al 2013), in this study, we investigated whether the connectivity varies depending on the expression of a given IEG.

The regional expression density of c-Fos, Npas4, Arc positive cells was cross-correlated across a group of animals to generate a correlated IEG-expression matrix (Fig. 8A-C, left). The connectivity networks were then visualized by applying threshold (|R|>0.7) to the correlation matrix (Fig. 8A-C, right). The connectivity per brain area, or the number of edges per node, was higher in RC than HC in the c-Fos based network (Fig. 8D). Also, the connectivity per effective node, or the number of edges per non-zero node, was higher in RC and in CFC than HC in the c-Fos and Npas4 based networks, respectively (Fig. 8E). Overall, the connectivity network tended to be more complex by CFC and RC compared to HC (Fig. 8A-E). The double-/triple-IEG-based networks showed a similar tendency (Fig. S21); the connectivity per brain area was larger in CFC in c-Fos^+^/Npas4^+^ and in RC Npas4^+^/Arc^+^ networks than HC (Fig. S21E), and the connectivity per effective node was larger in CFC in c-Fos^+^/Arc^+^ and Npas4^+^/Arc^+^ networks than HC (Fig. S21F). The significant increases in the average correlation were observed in Npas4 and c-Fos^+^/Arc^+^ based networks (Fig. S21G). Comparing the graphs obtained from each cell group, the networks were more dissimilar across each cell group in CFC and RC than HC (Fig. 8F, G). Thus, networks of IEG-based functional connectivity tended to become more complex after mice received learning-related stimuli, but they were not identical between the types of IEG. This suggests that different IEG expression provides different information on the area-area connectivity in the brain.

**Figure 8:**
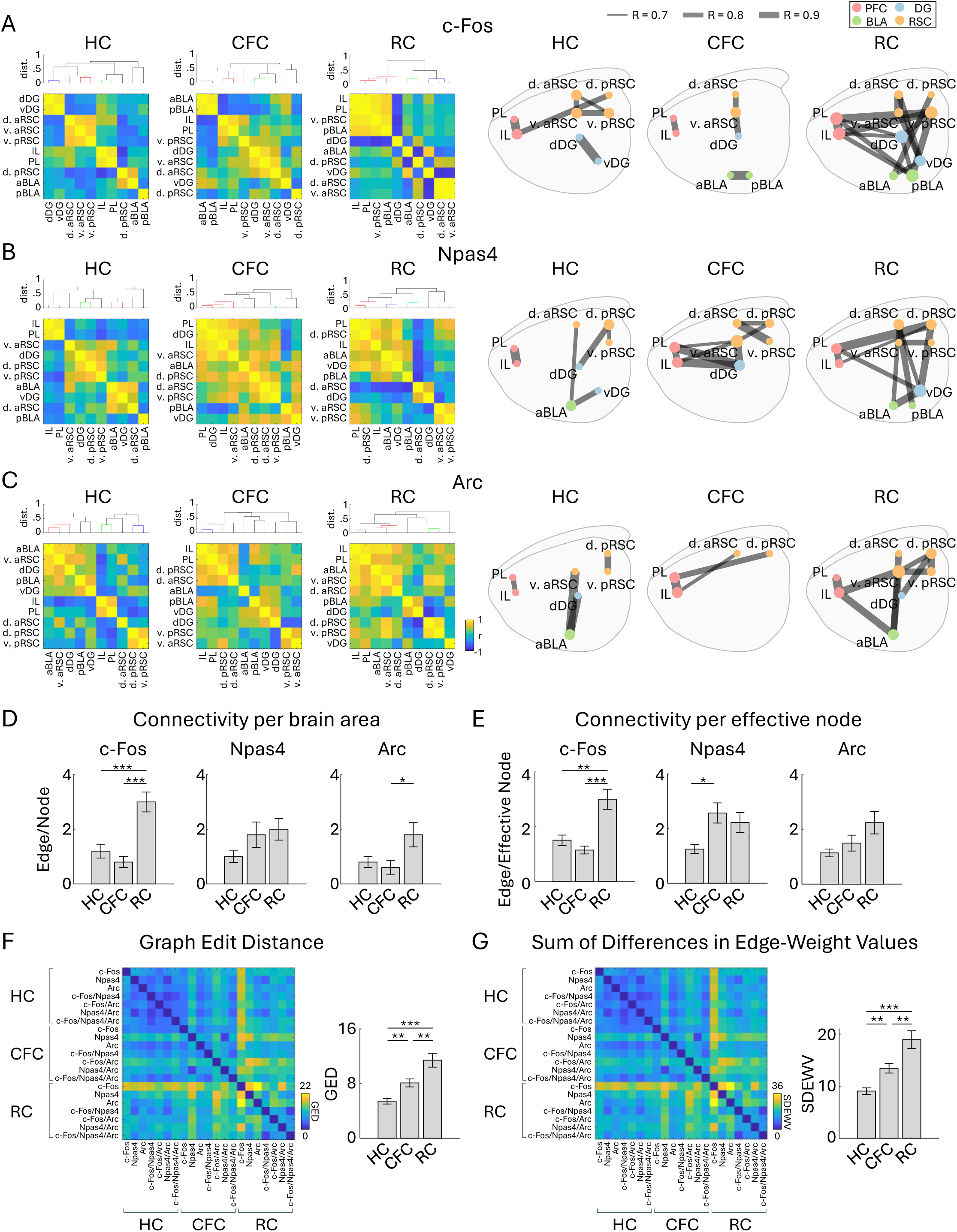
Functional connectivity network of each IEG. (A, B, C) Left, inter-regional correlation matrices for c-Fos- (A), Npas4- (B), and Arc- (C) positive cell densities. Dendrograms above the correlation matrices are calculated using dissimilarity index 1 - |r|, with colors indicating |r| > 0.7 (dissimilarity index < 0.3). Right, connectivity network graphs of c-Fos (A), Npas4 (B), and Arc (C) generated by connecting each brain region (node) based on the strong correlations (Pearson’s |r|L>L0.7) (right), for HC, CFC, and RC groups. The size of node circles in the network graphs corresponds to the number of connections (edges) the node has. (D, E) Quantification of network complexity. (D) Average number of edges per node across the ten brain regions. (E) Average number of edges per effective node, or the brain region which has at least one connection to another node, across brain regions. (F, G) Quantification of network dissimilarity. (F) Matrix of graph edit distance (GED) across the graphs of different IEG groups. Bar plot indicates average GED within HC, CFC, and RC groups. (G) Matrix of Sum of Differences in Edge-Weight Values (SDEWV) across the graphs of different IEG groups. Bar plot indicates average SDEWV within HC, CFC, and RC groups (n = 7 graphs for each).

## 4. Discussion

We found that expression patterns of c-Fos, Npas4, and Arc, along with their learning-induced changes, are not identical and vary depending on brain areas (Fig. 2, 3). The pattern of the combinative expression of those IEGs and their learning-induced changes also differ depending on brain areas (Fig. 4–7). We also demonstrate that different IEG expression provides different information on area-area connectivity networks (Fig. 8).

In this study, we used the method of automated cell detection after background assumption (ADABA) that we proposed previously (Fig. 1) (Osanai et al 2025). This method provides unbiased detection of the immunolabeled cells after reducing the effect of uneven background arising from biological structures in a brain tissue; for example, we often observed uneven background in the DG granule cell layer (Fig. S3, S4). Consistent with our previous report (Osanai et al 2025), our method provided highly precise cell detection, which aided the subsequent IEG co-expression analysis.

While several techniques for brain-wide neural activity mapping using IEGs have been developed in recent years (DeNardo et al 2019, Franceschini et al 2025, Guenthner et al 2013, Nagahama et al 2025, Renier et al 2016, Roy et al 2022, Wheeler et al 2013, Ye et al 2016, Zhang & Roy 2024), investigations of differences in IEG types are still limited (Chiaruttini et al 2024, Heroux et al 2018, Kawashima et al 2014, Ons et al 2004). Also, IEG expression in brain subregions has rarely been investigated comprehensively. In this study, we investigated c-Fos, Npas4, and Arc expressing cell densities in subregions of the PFC, BLA, DG, and RSC (Fig. 2, 3).

In the PFC, PL and IL subregions have distinct functions on fear expression/extinction and reward-seeking behaviors (Giustino & Maren 2015, Gourley & Taylor 2016, Sierra-Mercado et al 2011). The PL and IL have different c-Fos protein expression during fear renewal and retrieval of extinguished fear memories (Knapska & Maren 2009), and Arc RNA expression level is higher in the IL in the fear-extinguished rats but not different in the PL in the fear renewal (Orsini et al 2013). In contrast, both PL and IL show increased c-Fos protein expression after fear conditioning (Herry & Mons 2004) and cocaine conditioning (Zavala et al 2007) as well as increased Arc RNA expression after food conditioning (Schiltz et al 2007). In this study, we found that the number of c-Fos, Npas4 and Arc protein positive cells are similarly increased in both PL and IL in response to both fear and reward conditioning (Fig. 3; S7A, D).

In the BLA, c-Fos expression is induced by both aversive and appetitive stimuli, which are critical for valence-related behavior, observed using IHC and in situ hybridization (ISH) (Gore et al 2015, Redondo et al 2014). The aBLA and pBLA have neurons with different molecular identities and projections encoding negative and positive valence, respectively (Beyeler et al 2018, Kim et al 2016, O’Neill et al 2018, Pi et al 2020, Yang & Wang 2017, Zhang et al 2021, Zhang et al 2020). However, we did not observe such negative and positive valence differences in IEG expression changes between CFC and RC in the BLA although there were anterior-posterior differences in the type of IEGs whose expression increased; after both CFC and RC, c-Fos was significantly increased in the aBLA, whereas c-Fos, Npas4, Arc were increased in the pBLA (Fig. 3; S7G, J). Further cell-type-specific investigations (Zhang et al 2021, Zhang et al 2020) are needed to understand how valence-specific learning induces multiple IEG expression.

In the DG, dDG supports spatial memory while vDG is involved in fear and reward related behaviors (Kesner 2018, Kheirbek et al 2013, Kirk et al 2017). c-Fos in the DG has been shown to increase by conditioning to fear and reward in IHC (Beck & Fibiger 1995, Rademacher et al 2006), as well as Arc is increased by fear conditioning in ISH and transgene approach (Bal et al 2025, Rao-Ruiz et al 2019). Also, an increase of Npas4 has been observed with the RAM system (Sun et al 2020) and RNA level (Bal et al 2025). In this study, we observed an increase of c-Fos^+^ cells in both the dDG and vDG after fear and reward conditioning, and Arc increase in the dDG by reward and in the vDG by fear conditioning (Fig. 3; S10A, D). In contrast, we did not observe significant changes in the number of Npas4^+^ cells, which is similar to previous reports exploring Npas4 protein expression using fear conditioning (Chiaruttini et al 2024, Ramamoorthi et al 2011) and social interaction (Coutellier et al 2012). The discrepancy between our Npas4 expression results and the results from the RAM system and RNA level experiments implies that innate expression of Npas4 protein is strictly controlled unlike the RAM system or RNA level, but further investigations are needed.

Lastly, the RSC has several subregions with distinct cytoarchitectures, projections, and functions for episodic memory (Alexander et al 2023, Burwell & Amaral 1998, Cheng et al 2024, Sugar et al 2011, Sullivan et al 2023, Tsai et al 2022, Vann et al 2009, Vogt et al 2004). For example, the anterior part of the RSC is needed for object (de Landeta et al 2020) and trace-fear memory (Trask et al 2021), while the posterior RSC is important for spatial memory (de Landeta et al 2020, Trask et al 2021, Vann et al 2003). Dorsal (dysgranular) and ventral (granular) RSC are suggested to differentially encode allocentric and egocentric information (Alexander & Nitz 2015, Alexander et al 2023, Jacob et al 2017, Pothuizen et al 2009). IEGs including c-Fos and Arc are increased in the RSC after CFC in IHC (Robinson et al 2012). In subregional studies, c-Fos in both aRSC and pRSC is increased by acquisition of CFC memory and recall, whereas another IEG, zif268, is increased only in the aRSC during recall phase in IHC (Trask & Helmstetter 2022). The c-Fos expression in the both dorsal and ventral RSC is increased by fear conditioning in IHC (Radwanska et al 2010), but have differential expression in a spatial working memory task; c-Fos increase is observed both in the dorsal (dysgranular) and ventral (granular) RSC with visual cue, although more clearly in the posterior part, whereas the increase is observed only in the ventral RSC in the dark in IHC (Pothuizen et al 2009). Supporting subregional differences of the RSC, we found IEG expression differences in the RSC subregions (Fig. 3; S8E, G, I, K). For instance, while c-Fos was increased by CFC in the dorsal aRSC and in the dorsal/ventral pRSC, the increase was not significant in the ventral aRSC. Also, Npas4 was increased by RC in the dorsal/ventral aRSC but it was not clear in the dorsal/ventral pRSC. Our multi-region multi-IEG analysis supports that the induction of each IEG differs depending on the subregion and may contribute to the distinct memory functions.

We observed the brain region-dependent enhancement of combinative expression of cFos, Npas4, and Arc following aversive or rewarding experiences (Fig. 4–7). While there are several studies investigating colocalization of different IEGs (Chan et al 1993, Gonzales et al 2020, Guldenaar et al 1994, Guzowski et al 1999, Guzowski et al 2001, Hrvatin et al 2018, Lonergan et al 2010, Nakagami et al 2013, Sheng et al 1995, Stone et al 2011, Thompson et al 2010, Zuniga et al 2024), comprehensive analysis across brain areas have rarely been conducted. Recently, Chiaruttini et al. developed a pipeline to investigate brain-wide IEG expression and demonstrated the colocalization of c-Fos/Npas4 and c-Fos/Arc in HC, novel context, and CFC using IHC (Chiaruttini et al 2024). They observed basal co-expression level in HC varies across brain areas, with high abundance of c-Fos/Arc colocalization in the DG. They also observed co-expression of c-Fos/Arc increased in the DG, BLA, and some cortical areas by novel context exposure and CFC, which is consistent with our results. In DG area analysis, we found that the ratio of triple co-expressed neurons was different between the dorsal and ventral DG (Fig. 6, S11E, F). In addition, we found that the increase of c-Fos/Npas4/Arc triple-expressing cell ratio was less clear in d/vDG compared with in the PFC and in BLA unlike the increase of c-Fos/Arc double co-expression. This indicates that correlation of different expression varies depending on the combination of co-expression pattern of IEG types. Thus, although different types of IEGs tend to express in a neuron collaboratively (Fig. 4–7), the degree of co-expression might depend on the animal’s states, brain areas, and IEG types.

Do engram cells tagged by different IEGs or co-expressed IEGs play different roles in memory? By viral vector approaches, Sun et al. recently found c-Fos^+^ and Npas4^+^ cell populations labeled by RAM system in DG have distinct roles in memory generalization and discrimination (Sun et al 2020). Ye et al. found that Arc^+^/Npas4^+^ cells in PFC are involved in positive-valence experience but not entire Arc^+^ or Npas4^+^ populations as well as involvement of c-Fos^+^/Npas4^+^ cells than entire Fos^+^ population using Npas4-IHC and Arc-dependent TRAP mouse or fosCreER virus (Ye et al 2016). However, this question remains largely open. Although neural activity and IEG expression are strongly coupled, the expression does not simply reflect the level of average neural activity. During new context exploring, only fractions of CA1 place cells are tagged by c-Fos (Tanaka et al 2018). In hippocampal culture, c-Fos expression is induced by synchronized input activity with preference at 0.1 Hz and 50 Hz but not solely by raising cAMP, suggesting a relationship with sharp-wave-ripples and gamma oscillations (Anisimova et al 2023, Gee et al 2024, but see Yang et al 2024). In contrast, Arc transcription peaks with 10 Hz stimulation, suggesting a relationship with theta oscillations (Kim et al 2024). Also, the correlation between physiological neural activity and IEG expression is not constant across IEG types: using the FosGFP (Barth et al 2004) and EGFP-Arc mice (Okuno et al 2012), the correlation of neural calcium activity with c-Fos expression is higher than with Arc in the CA1 (Mahringer et al 2019) and visual cortex (Mahringer et al 2022). These suggest that IEG expression may reflect cellular functions including synaptic plasticity than merely indicating neural activity, while c-Fos and Arc are not always required for induction of long-term potentiation (Douglas et al 1988, Kyrke-Smith et al 2021, Wisden et al 1990). Given that different IEGs play different roles in synaptic plasticity, neurons expressing multiple IEGs could be influenced by the animal’s experience more than neurons expressing a single IEG, which may play an important role in contributing to diverse forms of synaptic plasticity in learning. In hippocampal culture, c-Fos^+^/Arc^+^ cells identified by IHC increase correlated cell firing following chemically induced long-term-potentiation, whereas c-Fos^-^/Arc^+^ cells decrease correlated cell firing, suggesting that different IEGs and their combination perform distinct functions (Jiang & VanDongen 2021). Neurons in CA1 with high c-Fos induction show higher correlated activities than neurons with low c-Fos induction during the spatial learning in the Fos-GFP mouse (Pettit et al 2022). Also, Arc positive cells are more likely to participate in sharp-wave-ripples than the negative cells in the CA1 acute slices of Arc-dVenus mice (Norimoto et al 2018). Conditional knockout of Scg2, the gene activated by c-Fos, lowers fast-gamma oscillation power and shifts the preferred theta phase of spikes in CA1 (Yap et al 2021). Correspondingly, the spikes of CA1 c-Fos^+^ cells occur during fast gamma events than c-Fos^-^ cells, and the preferred theta phase of theta-burst spikes of c-Fos^+^ cells differs from c-Fos^-^ spikes (Tanaka et al 2018). The IEG-expressing cell assembles form spatially defined clusters in the striatum in ISH (Gonzales et al 2020). Together, IEG expression and their combinations may reflect ongoing synaptic plasticity which leads to local neural activity synchrony, with the specific form depending on the types of expressing IEGs.

Finally, we demonstrated the functional connectivity network across brain regions based on different IEGs (Fig. 8). Since a subpopulation of neurons activated in memory acquisition overlaps with those activated in recall (Josselyn & Tonegawa 2020, Roy et al 2022), we considered the possibility that the area-to-area network between these subpopulations is critical for memory recall and such subnetwork can be visualized by the co-expression of IEGs. However, we did not observe such correspondence of subnetwork in the double/triple IEG-based networks, compared with the c-Fos, Npas4 and Arc single IEG-based networks (Fig. 8A–C, S21A–D). On the other hand, although our correlation analysis used relatively small sample size (n = 6 animals for each group), we found systematic tendencies for the IEG-based functional connectivity networks to become more complex and more dissimilar between IEGs. It is reported that similar region- and IEG type-dependency in IEG-expression correlation was observed between hippocampus, entorhinal cortex, and visual cortex, and between RNA levels of c-Fos, Arc and zif268 (Guzowski et al 2001). These suggest the possibility that the different factors of functional connectivity are coded by the expression of different IEG or their co-expression. Consistent with our hypothesis, whisker association training does not alter c-Fos expression in the barrel cortex of the FosGFP mouse, suggesting that the c-Fos^+^ cells in the sensory cortex can be involved in other functions (Lee et al 2021). However, although IEG-based functional connectivity between brain areas has often been estimated (Franceschini et al 2023, Silva et al 2019, Takeuchi et al 2022, Tanimizu et al 2017, Vetere et al 2017, Wheeler et al 2013), direct evidence which supports the link between physiological- and IEG-based- connectivity remains scarce and requires a more detailed interpretation of IEG-based networks, for example investigating whether a specific IEG-expressing neuron preferentially connects to neurons expressing the same type of IEG.

In this study, we aimed to evaluate the differential or concurrent expression across three IEGs, which are often used as indicators of neural activity and memory engram cells. CFC and RC stimulation enhanced those IEG expression and prompted their connectivity networks. On the other hand, our investigations have several limitations. Because the animals were exposed to a novel context during CFC and RC, the IEG expressing cells encode the novel environment in addition to aversive and reward stimulations. Other behavior paradigms, e.g., novel context exposure, are needed to dissociate context and emotional components when we discuss the change of IEGs based on pure fear or reward value. The intensity or type of unconditioned stimulus may also affect IEG expression because different IEGs have different transcription induction thresholds (Abraham et al 1993, Worley et al 1993). Also, co-expression pattern can differ by observation time: the protein levels of c-Fos and Arc peak at 60–90 minutes and Npas4 peaks at 30–60 minutes after stimulation, while mRNA level of c-Fos and Arc peak at 30 minutes and Npas4 peaks at 5 minutes (Guzowski et al 2001, Lonergan et al 2010, Ramamoorthi et al 2011, Skar et al 1994, Sun & Lin 2016). Time-sensitivity and dynamics of IEG combinative expression needs to be investigated. It is also possible that the degree of observed IEG co-expression rate varies by the sensitivity of antibody probes. While we investigated the co-expression patterns of IEGs in multiple brain areas in this study, it would be interesting to investigate whether cells expressing different IEGs in same subregion have different anatomical long-range projections because the transcriptions and neural projections can differ depending on emotional valence (Fuentes-Ramos & Barco 2024, Shpokayte et al 2022, Ye et al 2016). In addition, we did not examine layer-specificity of IEG co-expression in the current study. Recently, several tools were developed for automated brain atlas registration (Chiaruttini et al 2024, Franceschini et al 2025, Terstege et al 2022) but they do not usually deal with layer structures. Development of automated registration techniques based on cytoarchitectures to identify brain subregions and layer structures will accelerate understanding of brain-wide IEG-expression patterns. Moreover, cell-type dependencies for IEG-expression have been reported (Gonzales et al 2020, Hochgerner et al 2023, Jaeger et al 2018, Lucas et al 2008, Yang et al 2022). Single-cell transcriptomes will help to reveal brain-wide IEG co-expression patterns with cell-type specificity (Chen et al 2019, Hrvatin et al 2018, Jovic et al 2022, Moffitt et al 2018, Tyssowski et al 2018, Wu et al 2017, Yao et al 2023). However, it should be noted that the levels of RNA and proteins can mismatch due to complex posttranscriptional processes (Alberini & Kandel 2014, Buccitelli & Selbach 2020, Guzowski 2006, Li et al 2020). These suggest that the IEG-RNA expression is more relevant to neural activity, while protein expression is more relevant to cellular functions. Further studies are needed to investigate simultaneous recording of neural activity and dynamics of IEG RNA (Lee et al 2022) and protein synthesis (Meenakshi et al 2021), to understand whether RNA and protein tag engram cells similarly or not. Also, it is notable that IEG proteins have wide-range functions beyond affecting synaptic plasticity, including lipid synthesis (Caputto et al 2014, Rodriguez-Berdini et al 2020, Vaughen et al 2023), DNA repair (Pollina et al 2023), protection against neuronal death (Rawat et al 2016), and β-amyloid generation (Wu et al 2011). Given this background, it could be considered that various cellular functions can be interpreted from IEG-tagged engram cells.

In conclusion, we found that basal and learning-induced expression of c-Fos, Npas4, Arc, and their combinations vary across different brain areas. The results of IEG-based connectivity analysis suggest that different functional connectivity is coded by the expression of different IEG or their co-expression. These findings provide insights that engram cells also can be differently identified depending on the types and the combinations of IEGs. Further investigations are needed to understand whether interactions between different IEGs contribute unique roles in memory, in order to gain more detailed functional understanding and interpretation of IEG-tagged engram cells.

## Author contributions

H.O., T.K., and S.K.O. contributed to the study design. M.A. and H.O. conducted experiments. H.O. conducted all analysis, M.A conducted manual cell detection, and C.S. visualized network graph. H.O., T.K., and S.K.O. wrote the manuscript. All authors approved the final manuscript.

## Declaration of Competing Interest

The authors declare no competing interests.

## Data availability

Data will be made available on request. The custom automated cell detection MATLAB code used in this study is available at https://github.com/HisayukiOsanai/CellDetection.

## Acknowledgements

We thank all members of the Kitamura laboratory for their support. This work was supported by grants from the Endowed Scholar Program to T.K., the National Institute of Mental Health to T.K. (R01MH120134, R01MH125916), the National Institute of Neurological Disorders and Stroke to T.K. (R01NS138075), and Japan-U.S. Brain Research Cooperative Program: FY2023 to M.A.

## Supplementary Figure Legends

**Supp. Figure S1:**
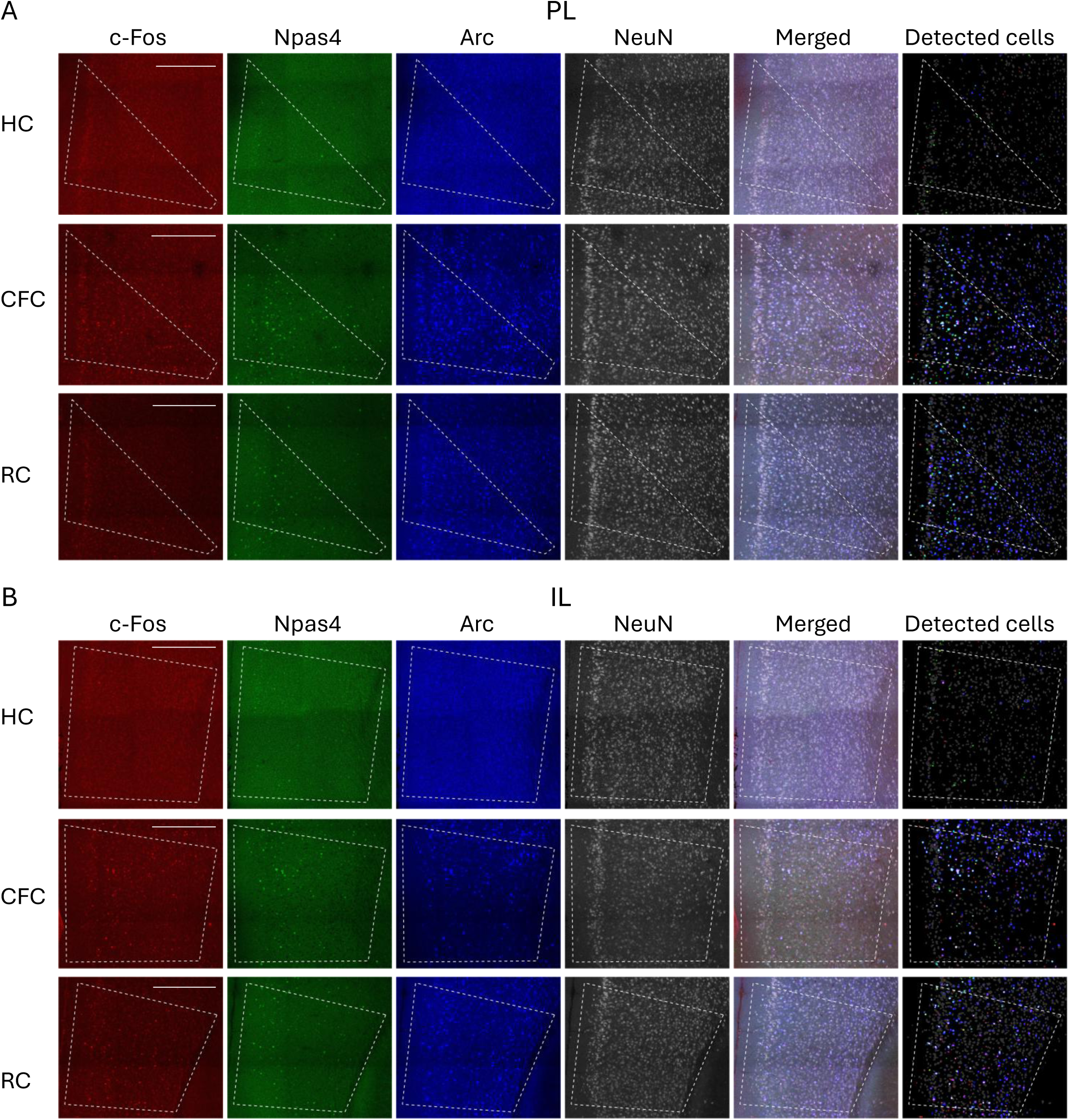
IEG expression in PFC (A, B) Larger field-of-view images of the PL (A) and IL (B). White dashed line indicates the region-of-interest (ROI) of each subregion used for automated cell detection analysis. Scale bars, 400 µm.

**Supp. Figure S2:**
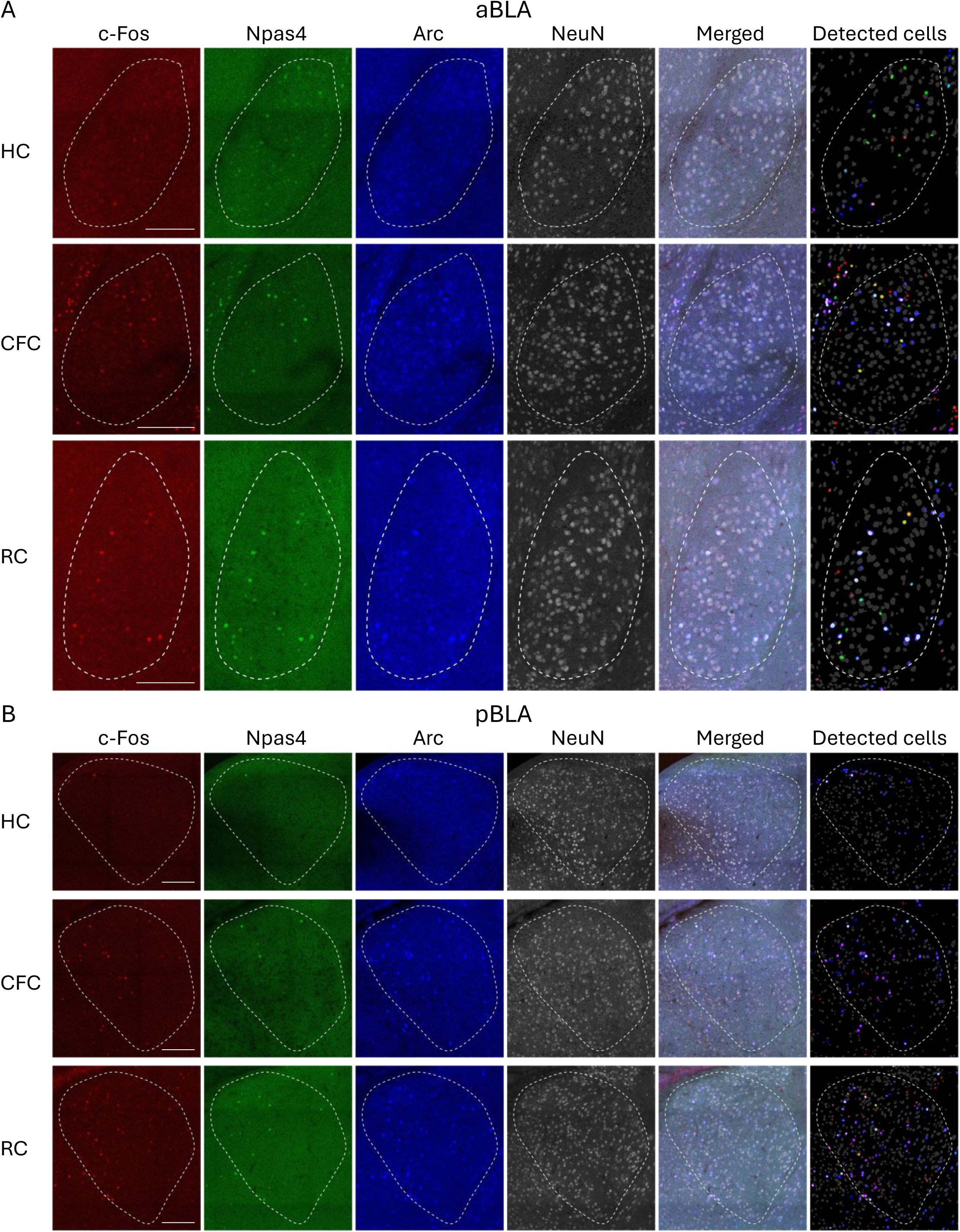
IEG expression in BLA (A, B) Larger field-of-view images of the aBLA (A) and pBLA (B). White dashed line indicates the ROI of each subregion. Scale bars, 200 µm.

**Supp. Figure S3:**
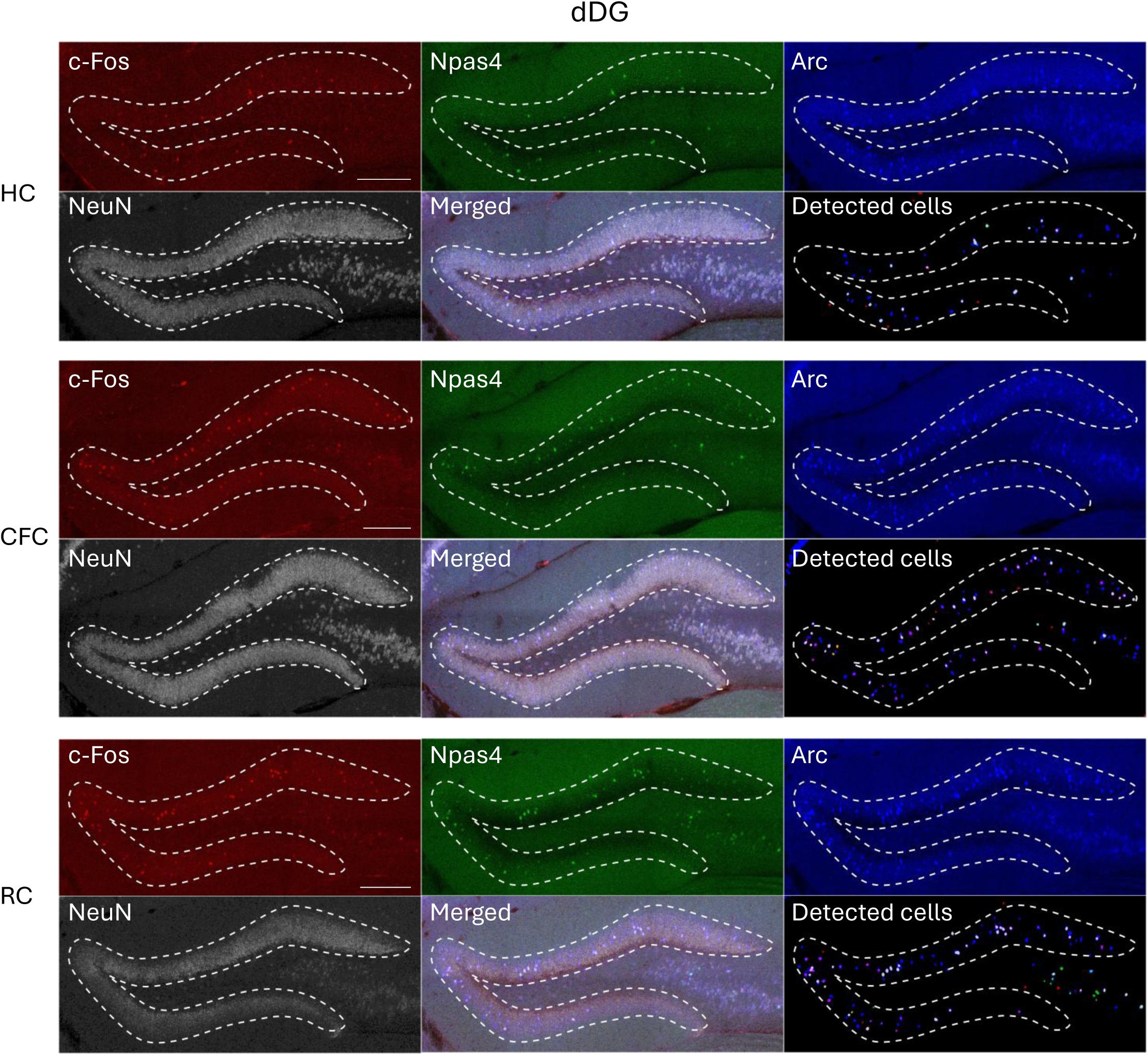
IEG expression in dDG Larger field-of-view images of the dDG. White dashed line indicates the ROI. Scale bars, 200 µm. Uneven background was observed as the darker background level around subgranular zone of the granule cell layer in the Npas4 and Arc images.

**Supp. Figure S4:**
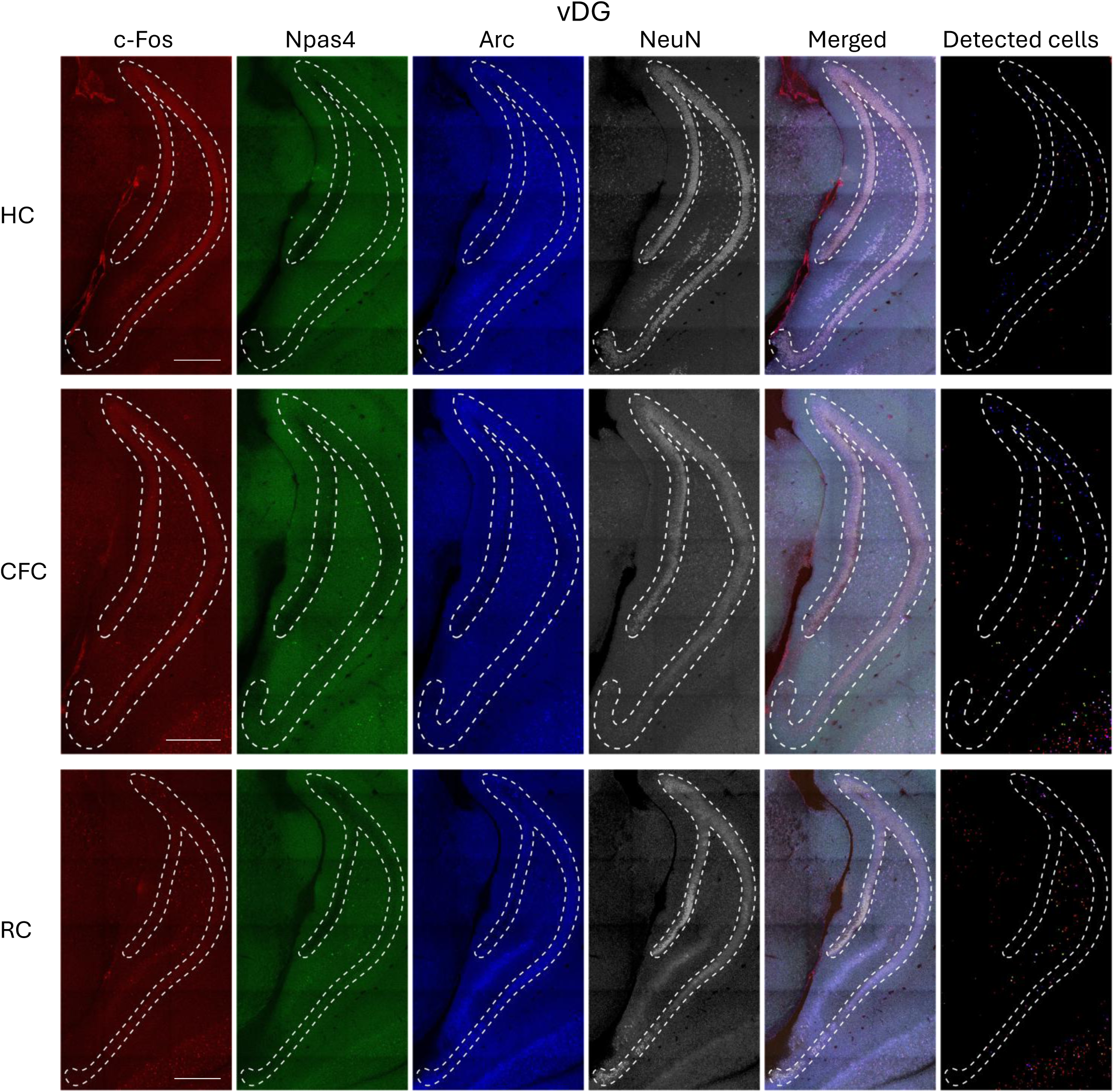
IEG expression in vDG Larger field-of-view images of the vDG. White dashed line indicates the ROI. Scale bars, 400 µm. Uneven background was observed in the c-Fos images of HC and CFC, and the Arc image of RC, as the increased autofluorescence along the granule cell layer.

**Supp. Figure S5:**
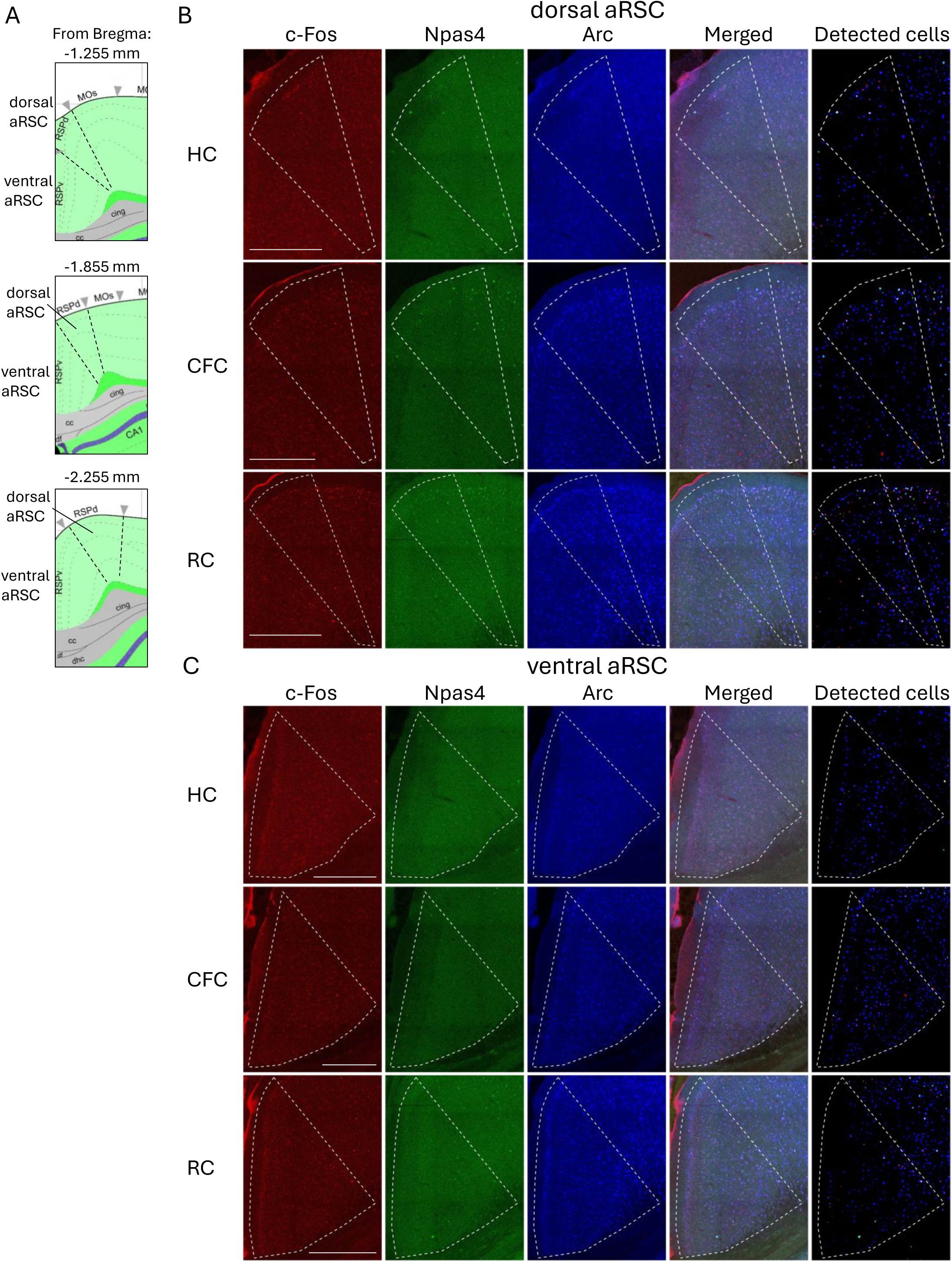
IEG expression in aRSC (A) Positions of the dorsal and ventral aRSC in the brain atlas (Allen Institute for Brain Science 2004). (B, C) Larger field-of-view images of the dorsal aRSC (B) and ventral aRSC (C). White dashed line indicates the ROI of each subregion. Scale bars, 400 µm.

**Supp. Figure S6:**
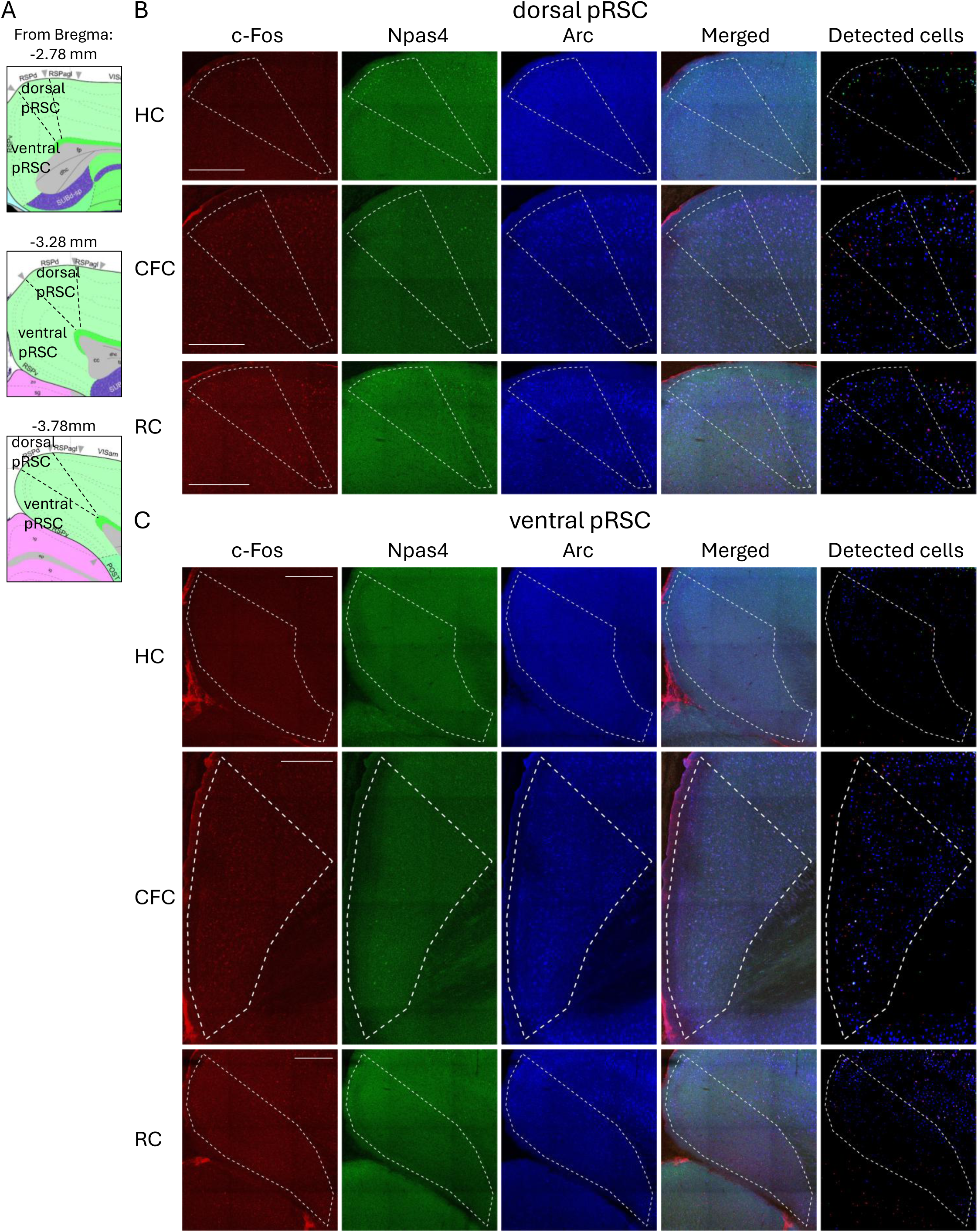
IEG expression in pRSC (A) Positions of the dorsal and ventral pRSC in the brain atlas (Allen Institute for Brain Science 2004). (B, C) Larger field-of-view images of the dorsal pRSC (B) and ventral pRSC (C). White dashed line indicates the ROI of each subregion. Scale bars, 400 µm.

**Supp. Figure S7:**
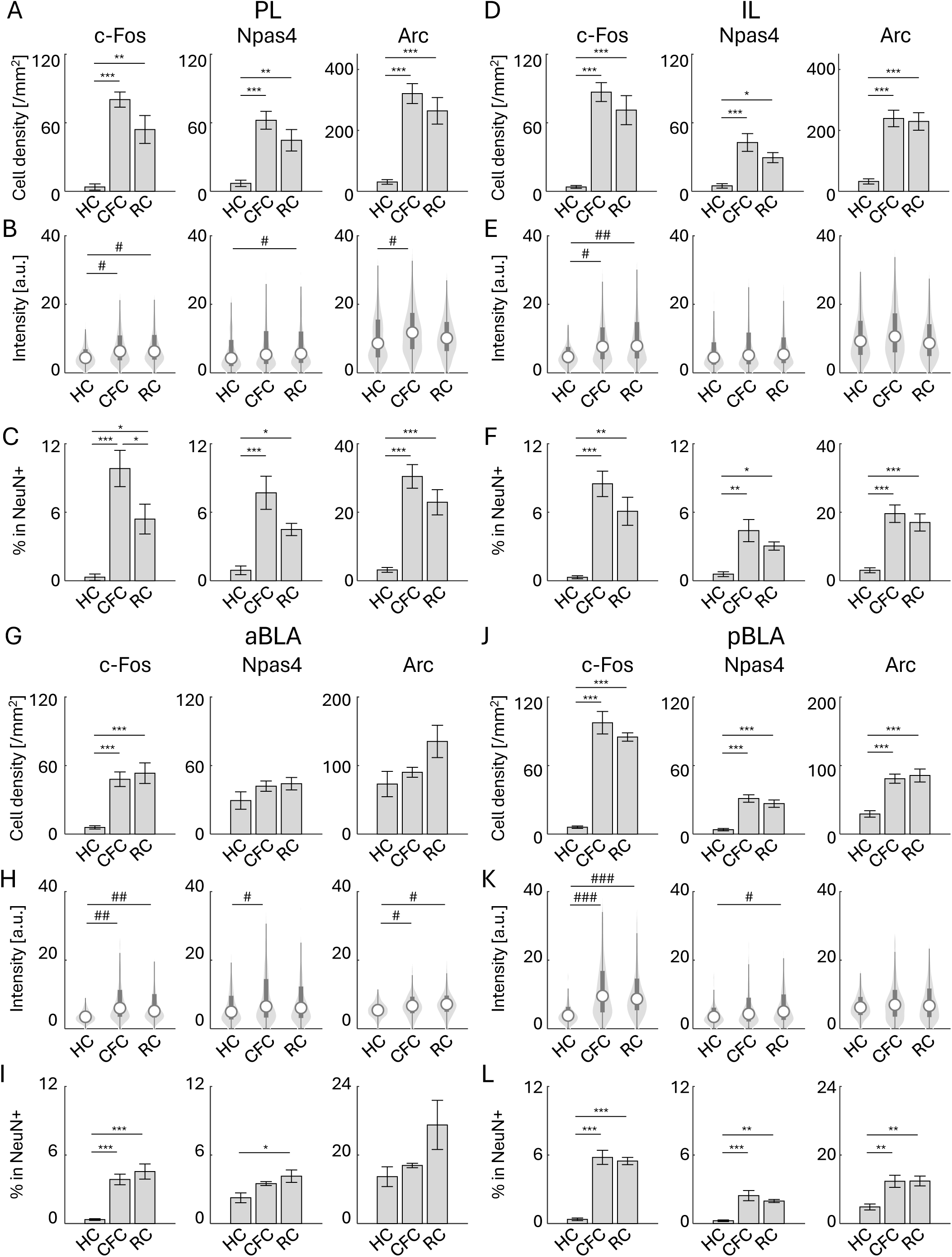
Cell density and expression level of IEG-positive cells in PFC and BLA (A–C), Analysis in the PL. (A) Cell density of c-Fos, Npas4, and Arc positive cells in HC, CFC, and RC in the PL. (B) Expression level of c-Fos, Npas4, and Arc positive cells in HC, CFC, and RC. n = 962, 4462, and 3897 cells for HC, CFC, and RC. (C) Percentage of c-Fos, Npas4, and Arc positive neurons per all NeuN^+^ cells. (D–F), Analysis in the IL. (D) Cell density of c-Fos, Npas4, and Arc positive cells in HC, CFC, and RC in IL. (E) Expression level of c-Fos, Npas4, and Arc positive cells in HC, CFC, and RC. n = 1228, 5058, and 4935 cells for HC, CFC, and RC. (F) Percentage of c-Fos, Npas4, and Arc positive neurons per all NeuN^+^ cells. (G–I), Analysis in the aBLA. (G) Cell density of c-Fos, Npas4, and Arc positive cells in HC, CFC, and RC in the aBLA. (H) Expression level of c-Fos, Npas4, and Arc positive cells in HC, CFC, and RC. n = 585, 987, and 1031 cells for HC, CFC, and RC. (I) Percentage of c-Fos, Npas4, and Arc positive neurons per all NeuN^+^ cells. (J–L), Analysis in the pBLA. (J) Cell density of c-Fos, Npas4, and Arc positive cells in HC, CFC, and RC in pBLA. (K) Expression level of c-Fos, Npas4, and Arc positive cells in HC, CFC, and RC. n = 437, 1110, and 1133 cells for HC, CFC, and RC. (L) Percentage of c-Fos, Npas4, and Arc positive neurons per all NeuN^+^ cells.

**Supp. Figure S8:**
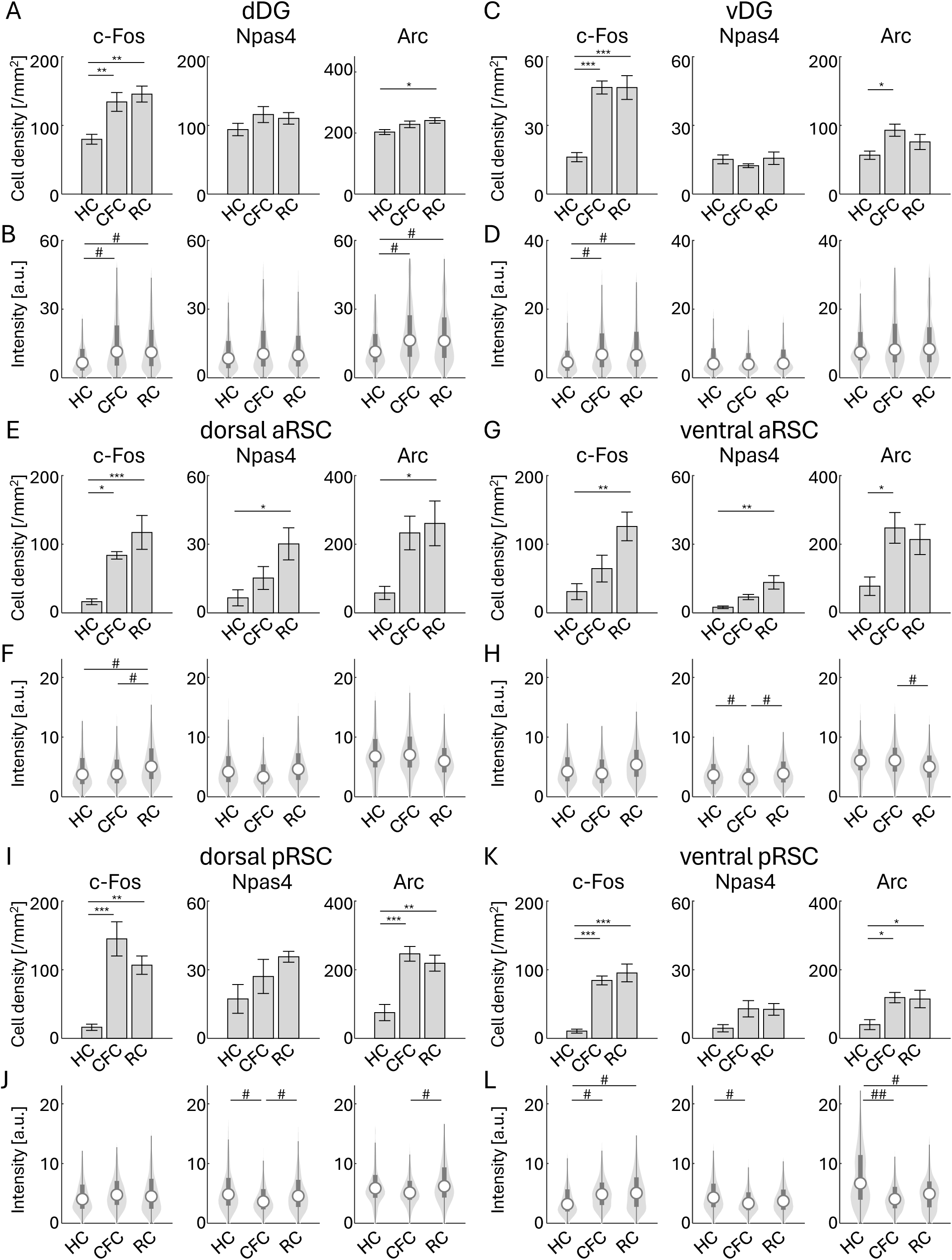
Cell density and expression of IEG-positive cells in DG and RSC (A, B), Analysis in the dDG. (A) Cell density of c-Fos, Npas4, and Arc positive cells in HC, CFC, and RC in the dDG. (B) Expression level of c-Fos, Npas4, and Arc positive cells in HC, CFC, and RC. n = 1413, 1542, and 1628 cells for HC, CFC, and RC. (C, D), Analysis in the vDG. (C) Cell density of c-Fos, Npas4, and Arc positive cells in HC, CFC, and RC in vDG. (D) Expression level of c-Fos, Npas4, and Arc positive cells in HC, CFC, and RC. n = 749, 1406, and 986 cells for HC, CFC, and RC. (E, F), Analysis in the dorsal aRSC. (E) Cell density of c-Fos, Npas4, and Arc positive cells in HC, CFC, and RC in the dorsal aRSC. (F) Expression level of c-Fos, Npas4, and Arc positive cells in HC, CFC, and RC. n = 783, 2103, and 2552 cells for HC, CFC, and RC. (G, H), Analysis in the ventral aRSC. (G) Cell density of c-Fos, Npas4, and Arc positive cells in HC, CFC, and RC in the ventral aRSC. (H) Expression level of c-Fos, Npas4, and Arc positive cells in HC, CFC, and RC. n = 1276, 2888, and 3068 cells for HC, CFC, and RC. (I, J), Analysis in dorsal pRSC. (I) Cell density of c-Fos, Npas4, and Arc positive cells in HC, CFC, and RC in the dorsal pRSC. (J) Expression level of c-Fos, Npas4, and Arc positive cells in HC, CFC, and RC. n = 1458, 2724, and 2102 cells for HC, CFC, and RC. (K, L), Analysis in the ventral pRSC. (K) Cell density of c-Fos, Npas4, and Arc positive cells in HC, CFC, and RC in the ventral pRSC. (L) Expression level of c-Fos, Npas4, and Arc positive cells in HC, CFC, and RC. n = 1955, 3931, and 4149 cells for HC, CFC, and RC.

**Supp. Figure S9:**
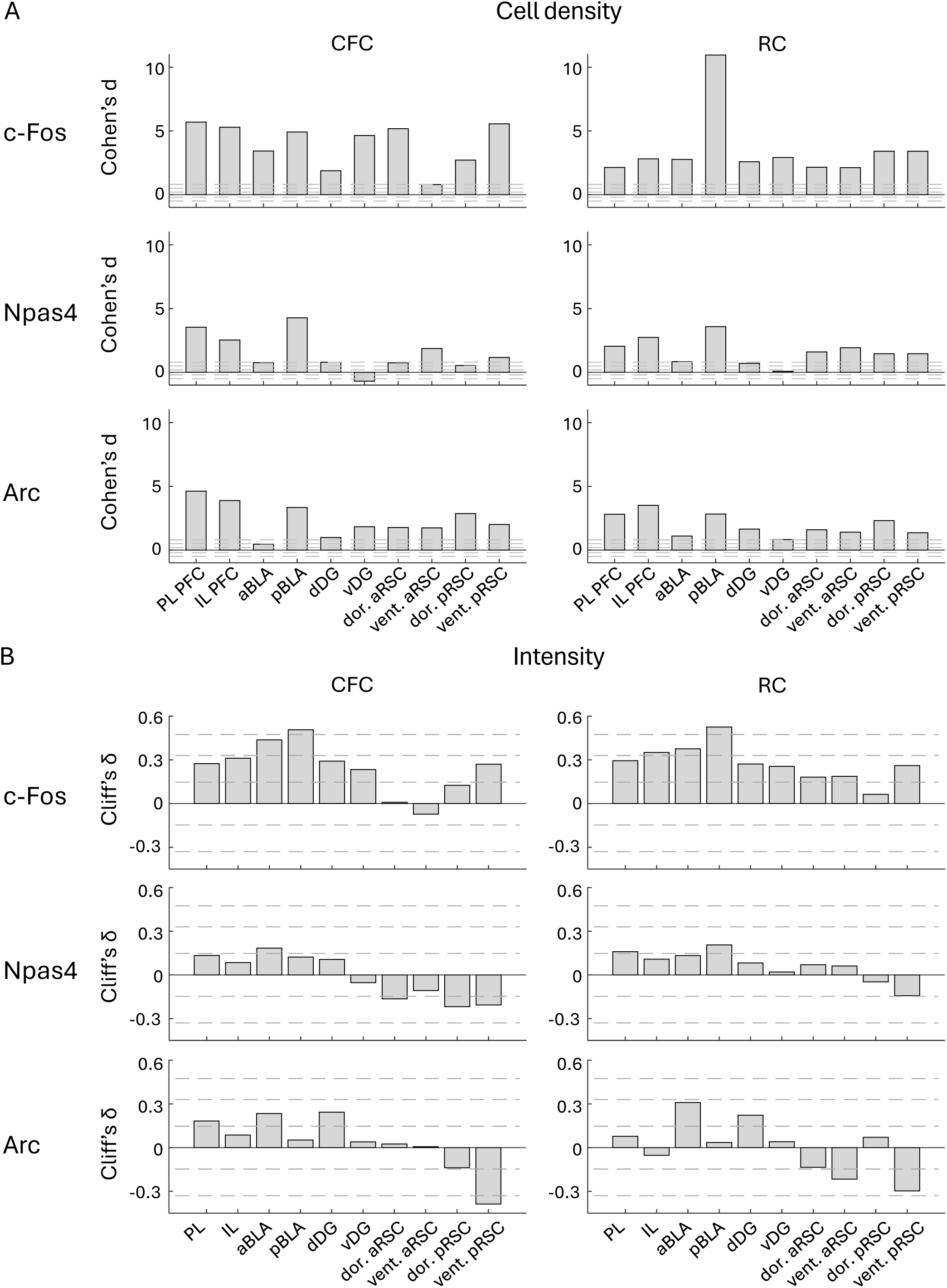
Effect size of cell density and intensity (A) Bar plots of Cohen’s d of cell densities across brain regions, calculated from the data shown in Supp. Figure S7A, D, G, J and S8A, C, E, G, I, K. Gray dashed lines indicate the effects are small (d = ±0.2), medium (d = ±0.5), and large (d = ±0.8). (B) Bar plots of Cliff’s delta of IEG intensities across brain regions, calculated from the data shown in Supp. Figure S7B, E, H, K and S8B, D, F, H, J, L. Gray dashed lines indicate the effects are small (δ = ±0.147), medium (δ = ±0.33), and large (δ = ±0.474).

**Supp. Figure S10:**
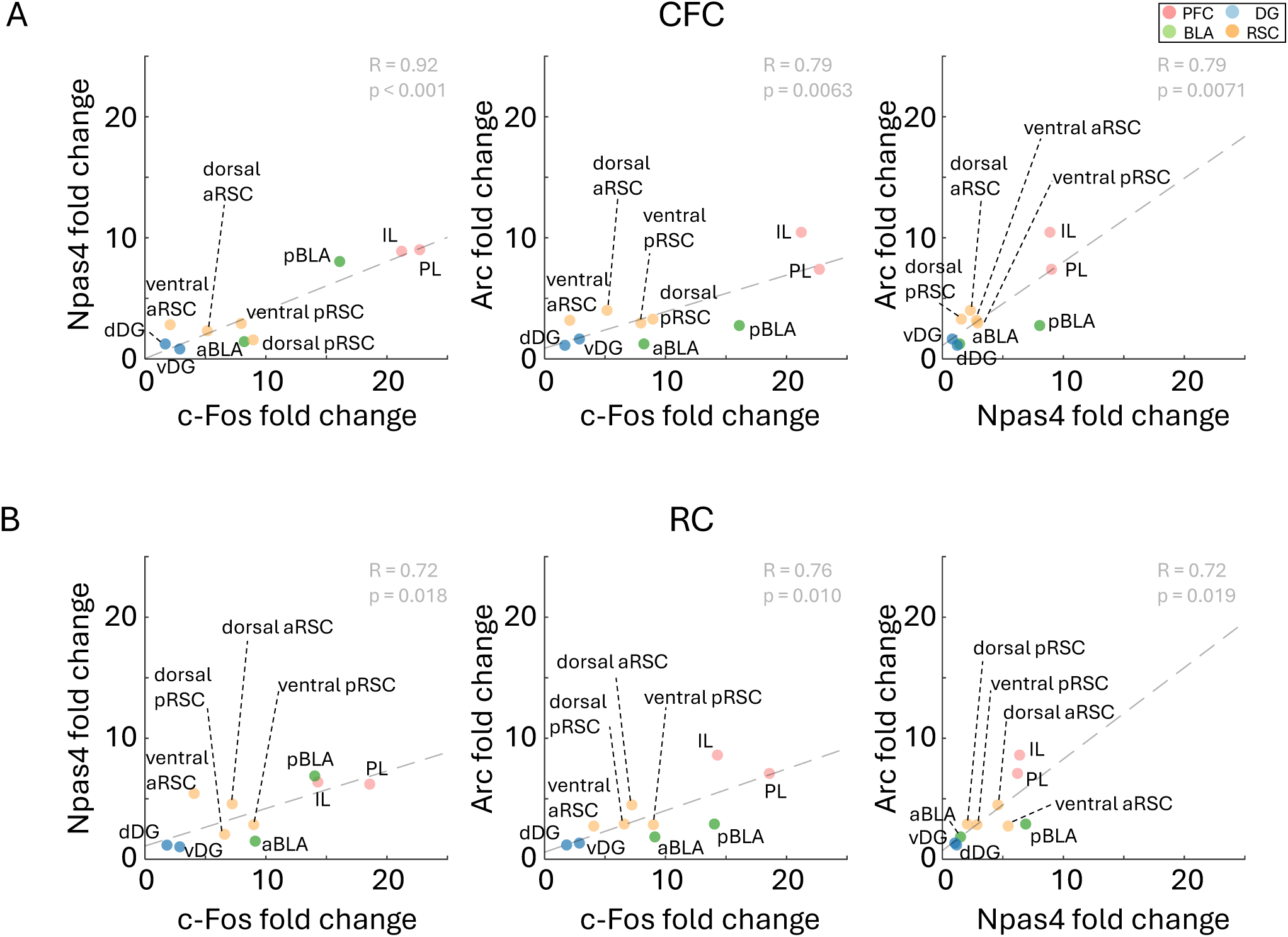
Cell density changes in each IEG in different brain regions (A) Scatter plots of fold-changes of c-Fos, Npas4, and Arc positive cell densities by CFC across ten brain regions, obtained from Figure 3A. (B) Similarly, scatter plots of fold-changes of c-Fos, Npas4, and Arc positive cell densities by RC, obtained from Figure 3A. Gray dashed lines indicate linear regression line. R and p indicate values of Pearson correlation.

**Supp. Figure S11:**
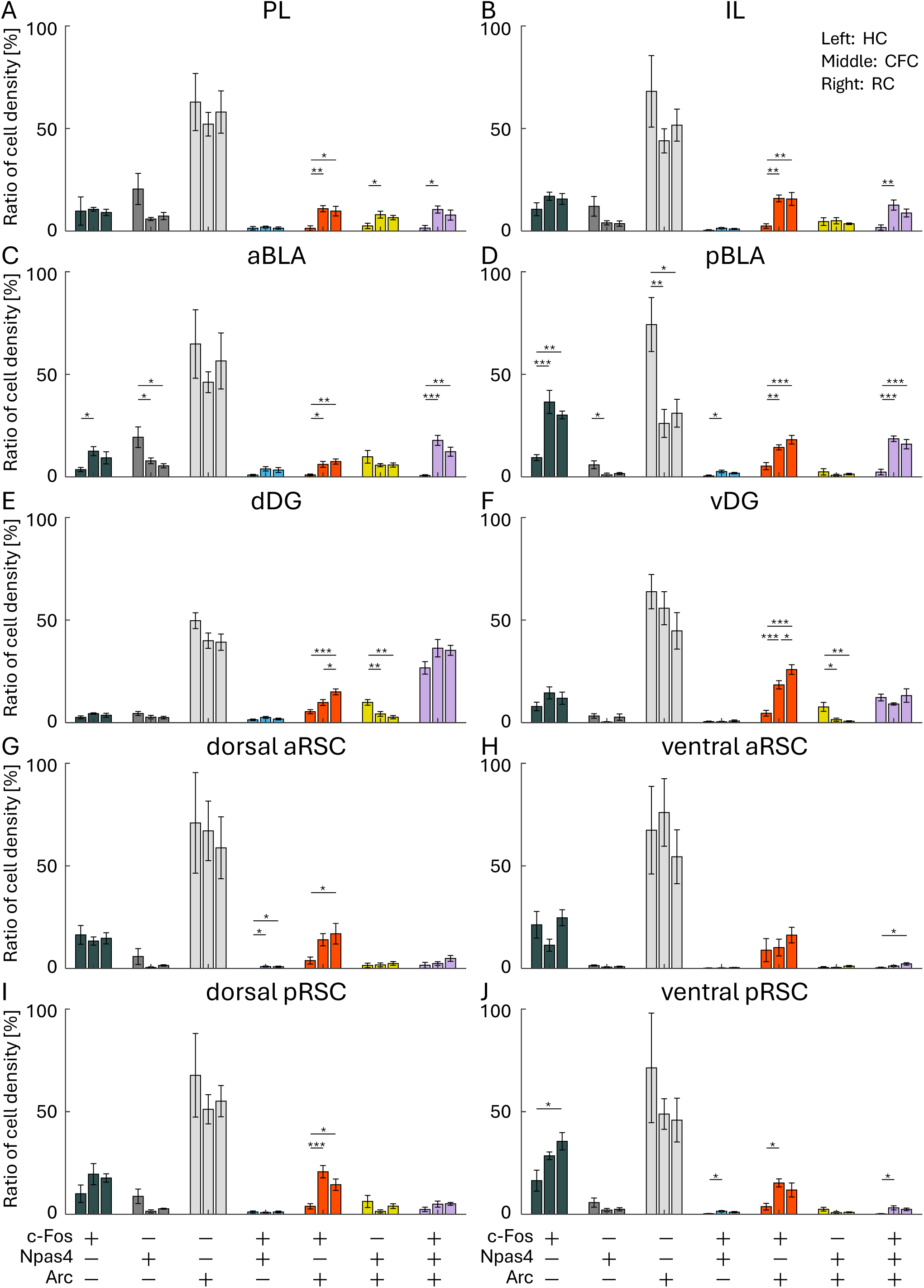
Cell density ratio per all IEG positive cells in each cell group Ratio of cell densities per all IEG positive cells. For each group of bars, the left bar indicates HC, the middle bar indicates CFC, and the right bar indicates RC groups. (A) PL, (B) IL, (C) aBLA, (D) pBLA, (E) dDG, (F) vDG, (G) dorsal aRSC, (H) ventral aRSC, (I) dorsal pRSC, and (J) ventral pRSC.

**Supp. Figure S12:**
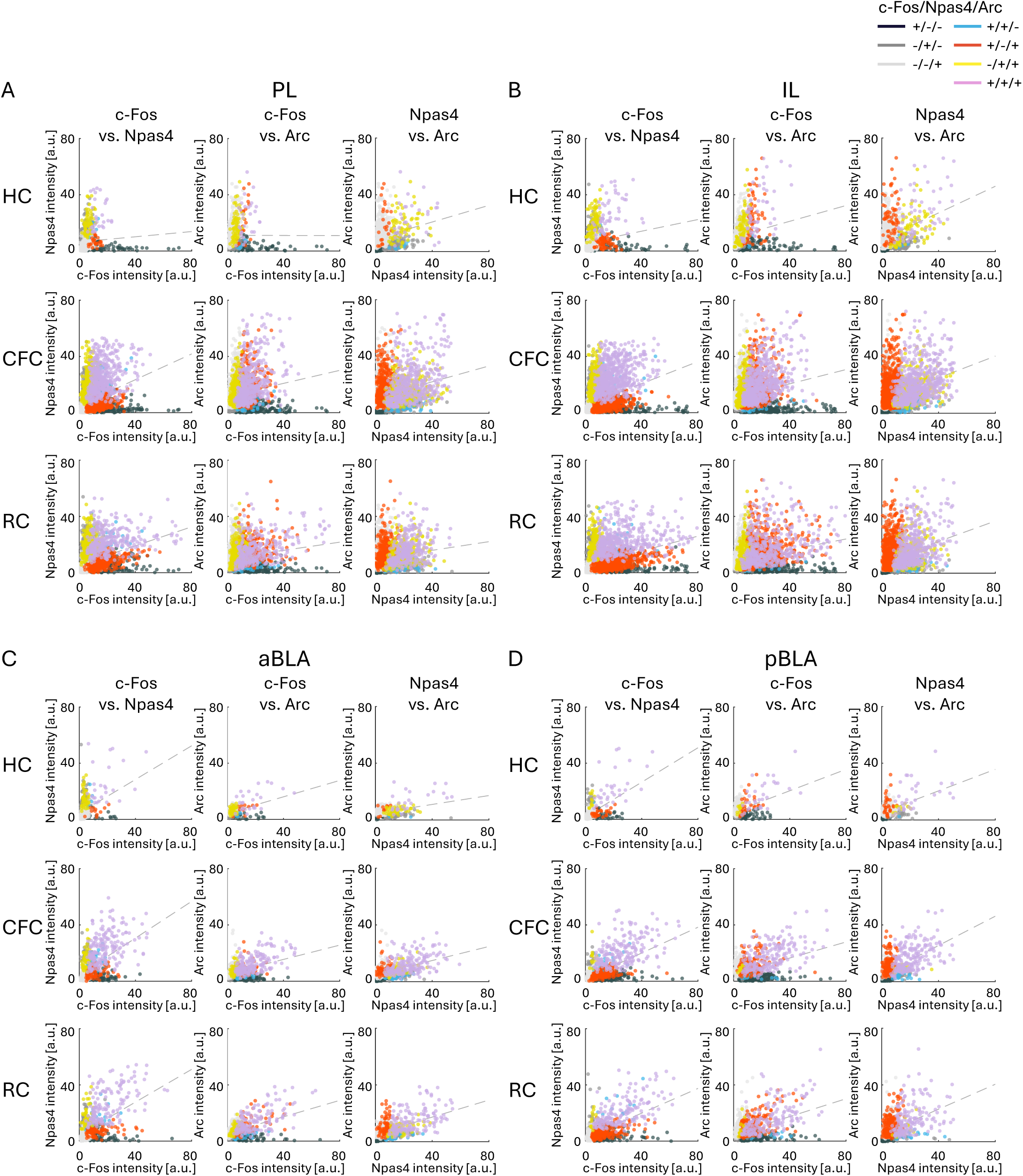
Intensities of IEGs in individual cells in PFC and BLA Scatter plots showing the intensities of c-Fos, Npas4, and Arc in single cells. (A) PL, (B) IL, (C) aBLA, and (D) pBLA. Colors represent cells with selective or concurrent expression of c-Fos, Npas4, and Arc. Gray dashed lines indicate correlations of cells across all groups. Intensities within each cell group are shown in Supp. Fig. S15. Correlation coefficients in each cell group are shown in Supp. Fig. S18.

**Supp. Figure S13:**
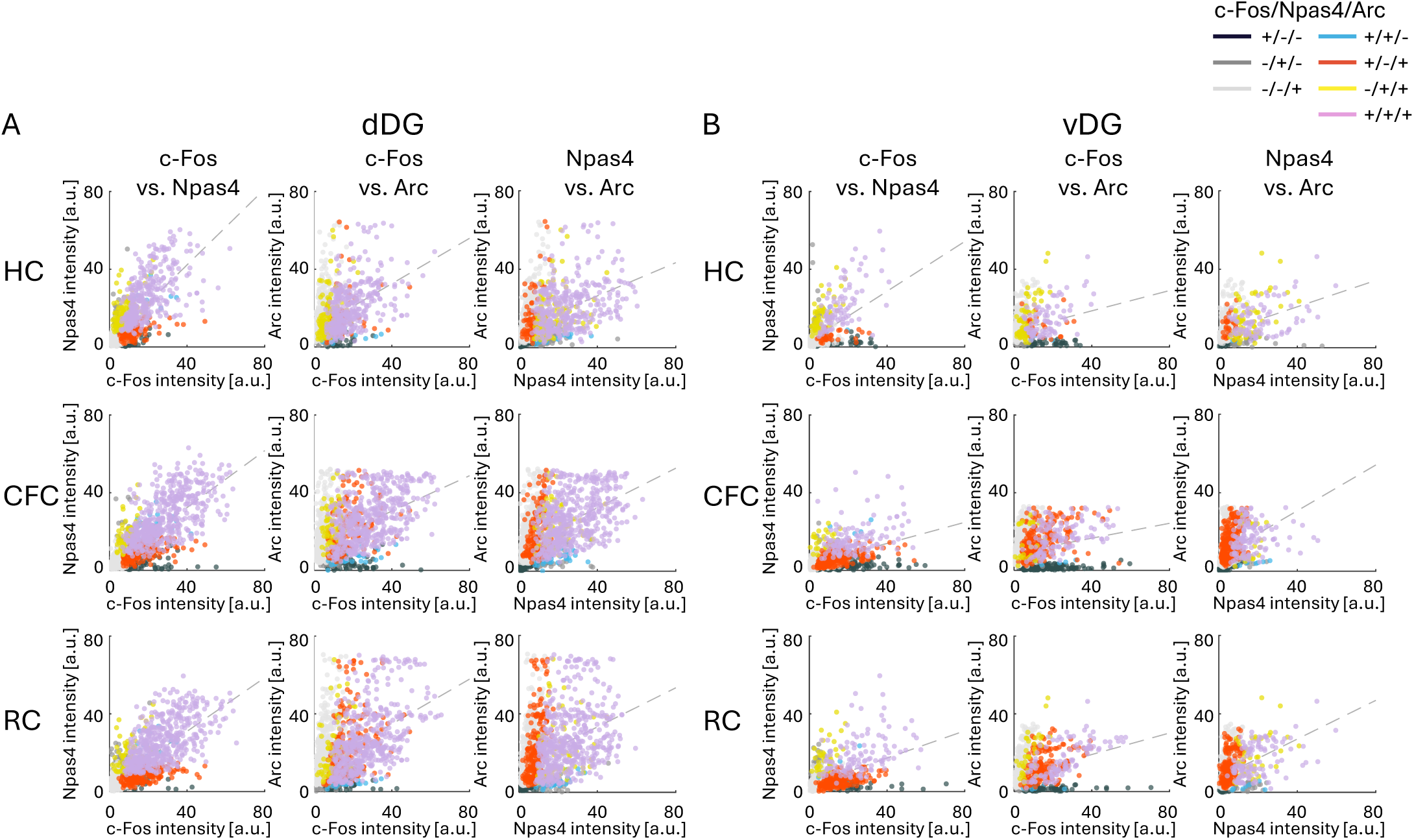
Intensities of IEGs in individual cells in DG Scatter plots showing the intensities of c-Fos, Npas4, and Arc in single cells. (A) dDG and (B) vDG. Colors represent cells with selective or concurrent expression of c-Fos, Npas4, and Arc. Gray dashed lines indicate correlations of cells across all groups. Intensities within each cell group are shown in Supp. Fig. S16. Correlation coefficients in each cell group is shown in Supp. Fig. S19.

**Supp. Figure S14:**
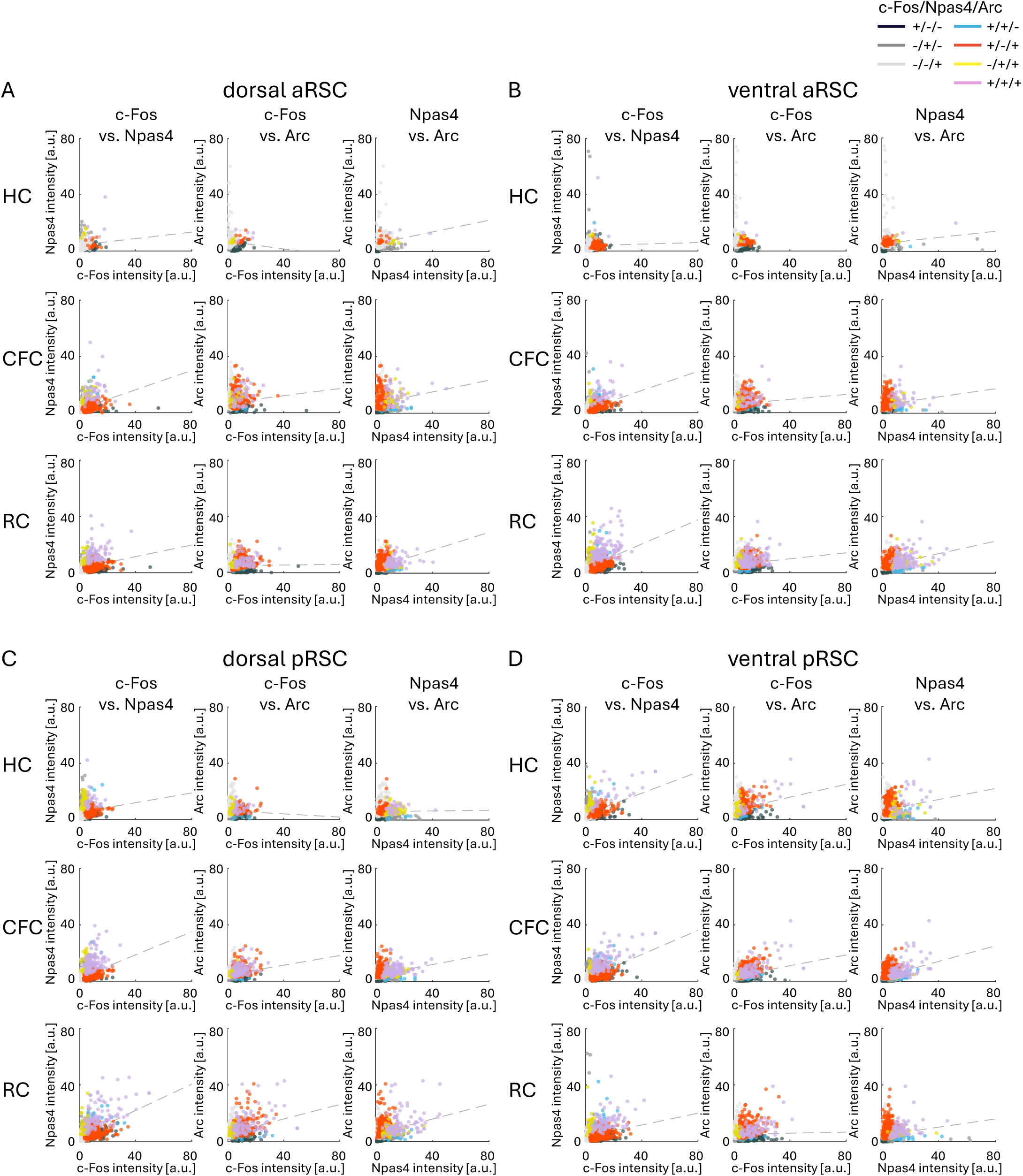
Intensities of IEGs in individual cells in RSC Scatter plots showing the intensities of c-Fos, Npas4, and Arc in single cells. (A) dorsal aRSC, (B) ventral aRSC, (C) dorsal pRSC, and (D) ventral pRSC. Colors represent cells with selective or concurrent expression of c-Fos, Npas4, and Arc. Gray dashed lines indicate correlations of cells across all groups. Intensities within each cell group are shown in Supp. Fig. S17. Correlation coefficients in each cell group are shown in Supp. Fig. S20.

**Supp. Figure. S15:**
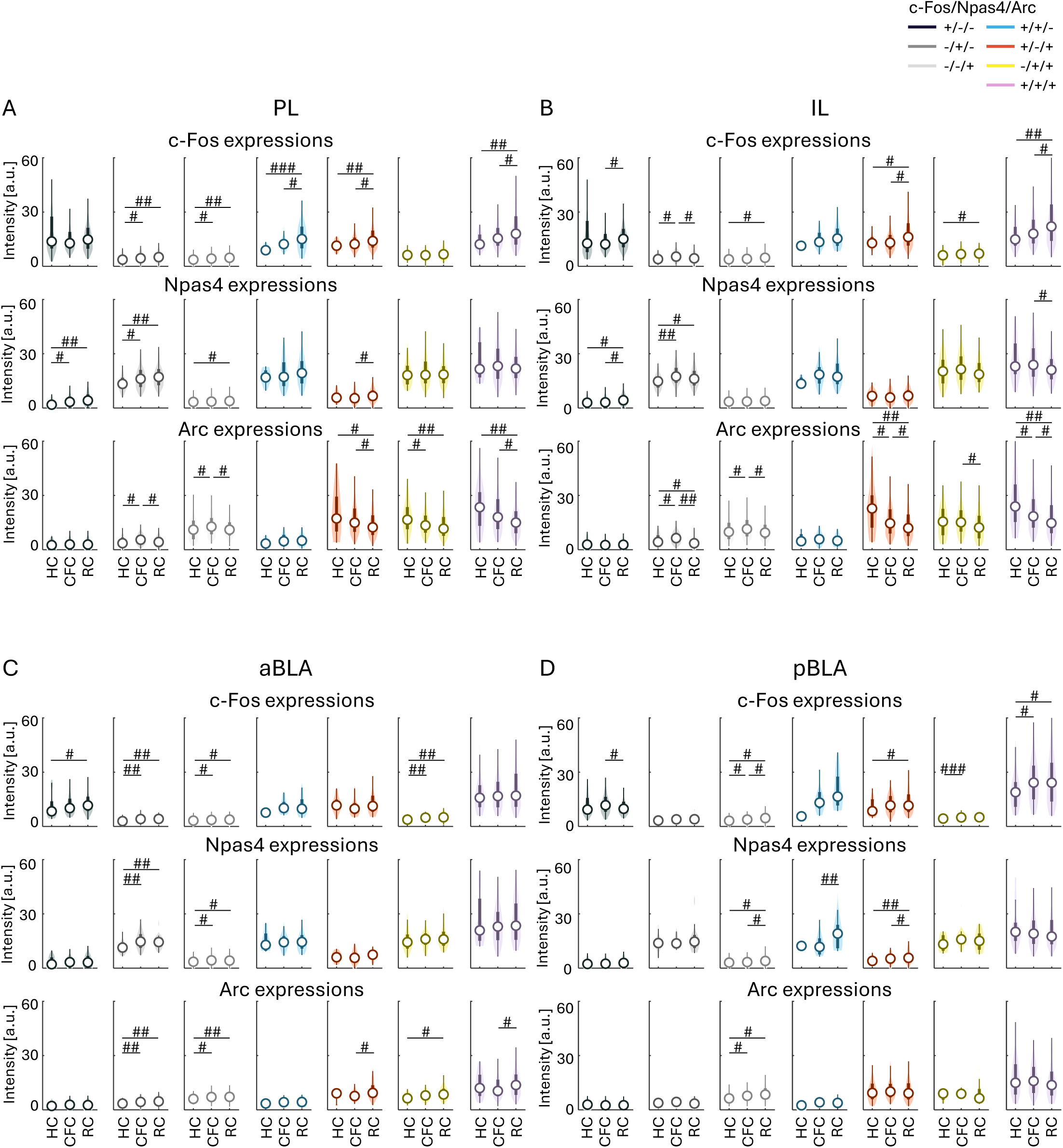
Intensities of IEGs in each cell group in PFC and BLA Intensities of c-Fos, Npas4, and Arc in individual cells in HC, CFC, and RC, in the PL (A), IL (B), aBLA (C), and pBLA (D).

**Supp. Figure S16:**
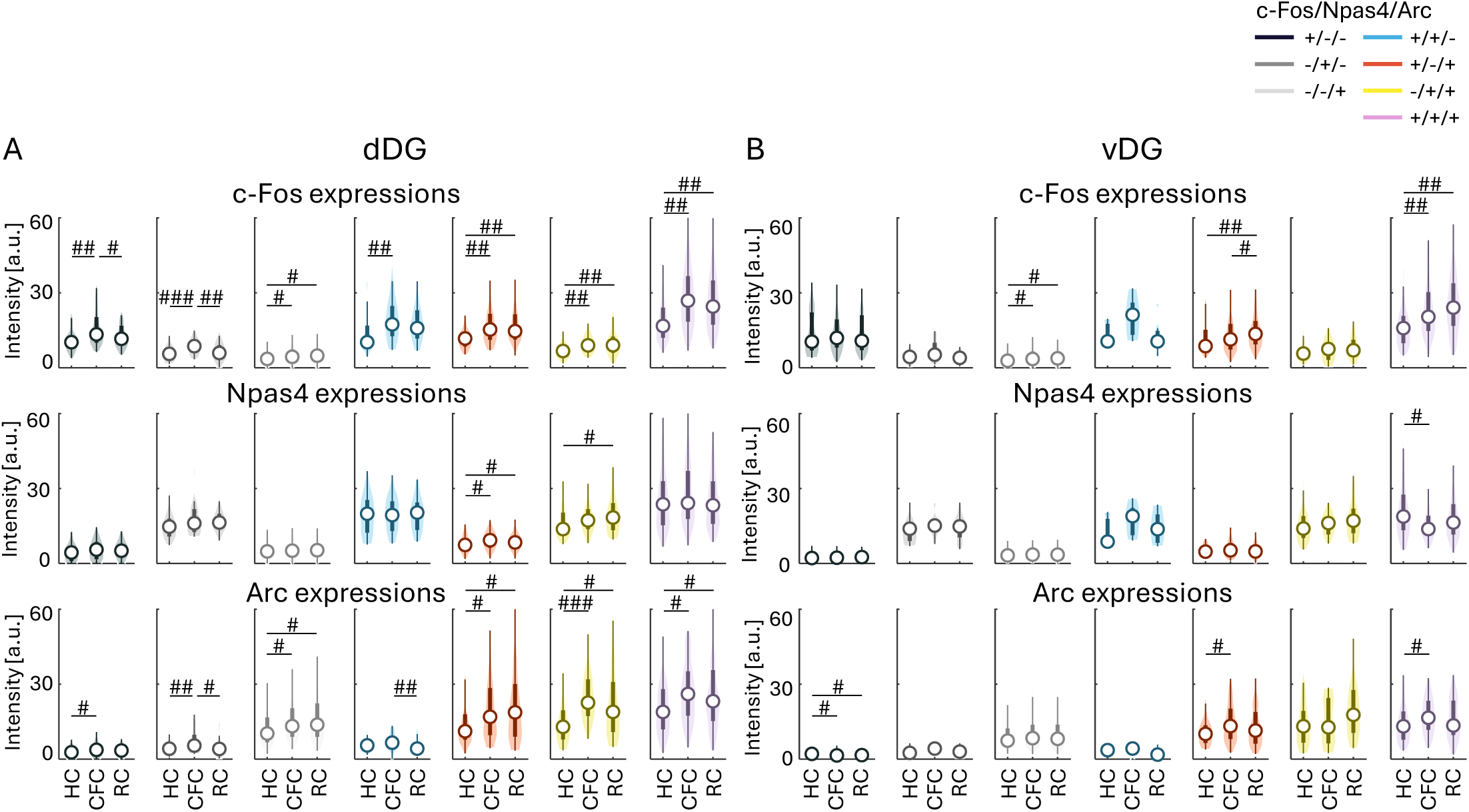
Intensities of IEGs in each cell group in DG Intensities of c-Fos, Npas4, and Arc in individual cells in HC, CFC, and RC, in the dDG (A) and vDG (B).

**Supp. Figure S17:**
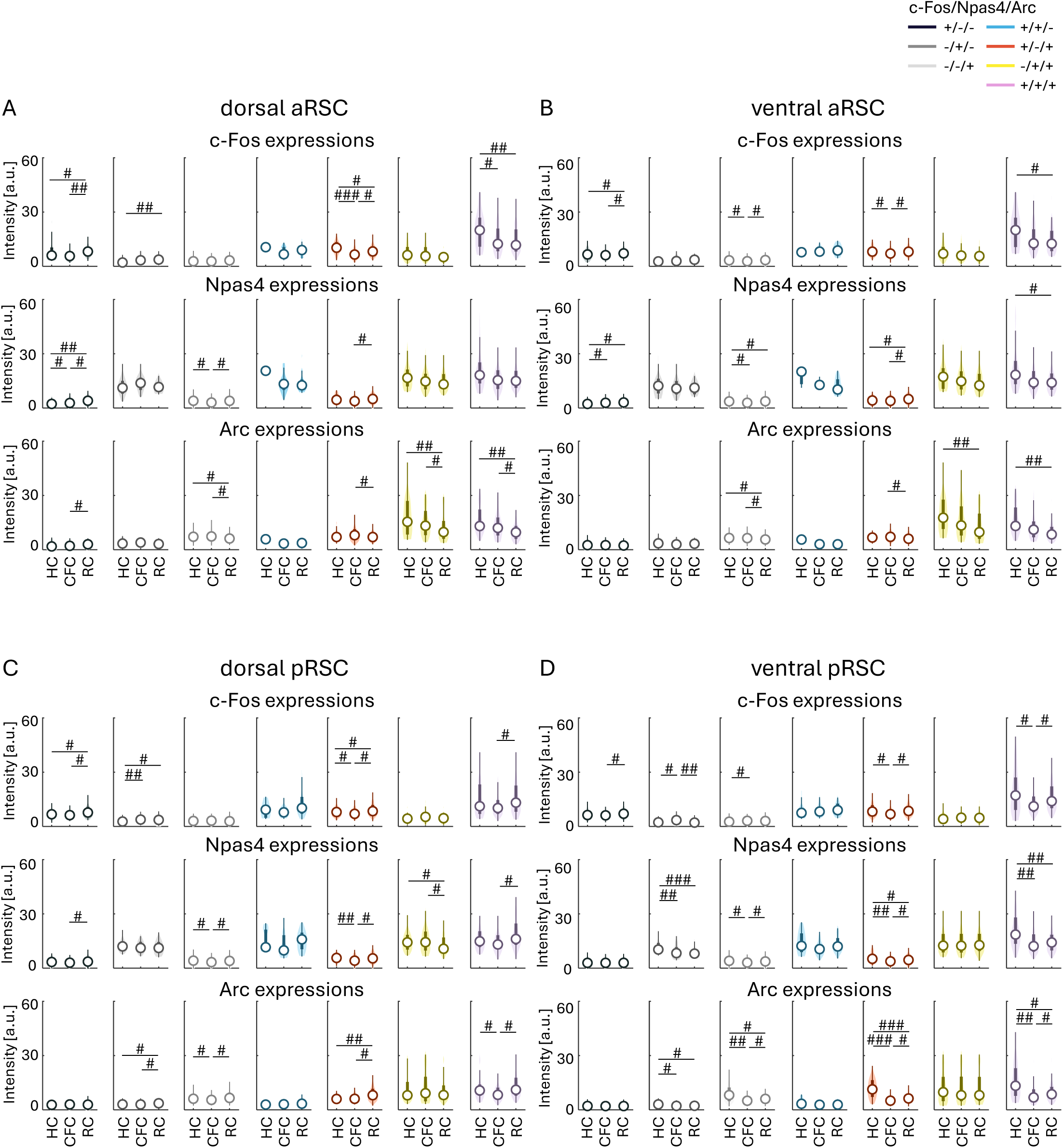
Intensities of IEGs in each cell group in RSC Intensities of c-Fos, Npas4, and Arc in individual cells in HC, CFC, and RC, in the dorsal aRSC (A), ventral aRSC (B), dorsal pRSC (C), and ventral pRSC (D).

**Supp. Figure S18:**
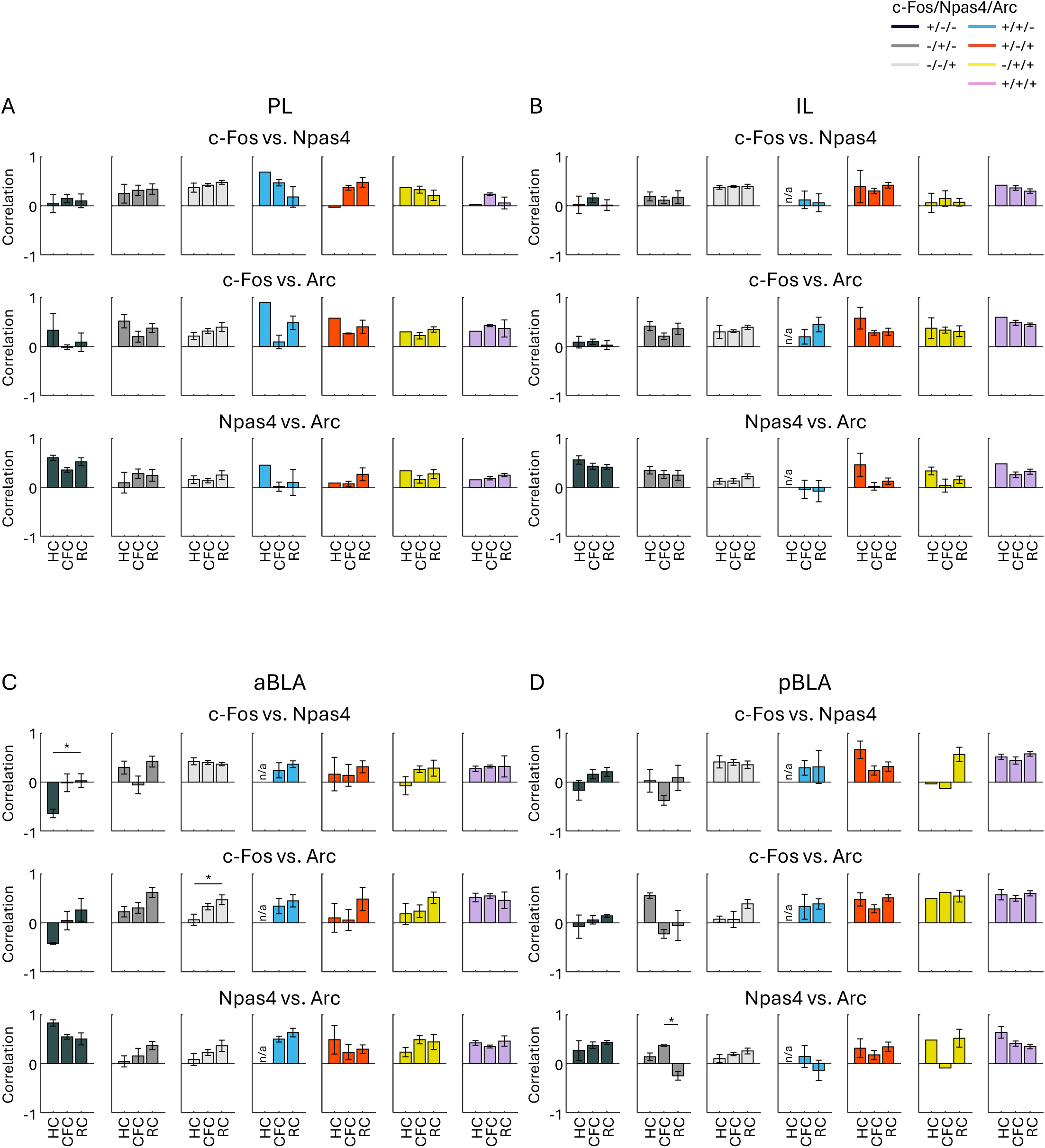
Correlation of IEG Intensities in each cell group in PFC and BLA Average correlation of IEG intensities of c-Fos, Npas4, and Arc in individual cells in HC, CFC, and RC, in the PL (A), IL (B), aBLA (C), and pBLA (D).

**Supp. Figure S19:**
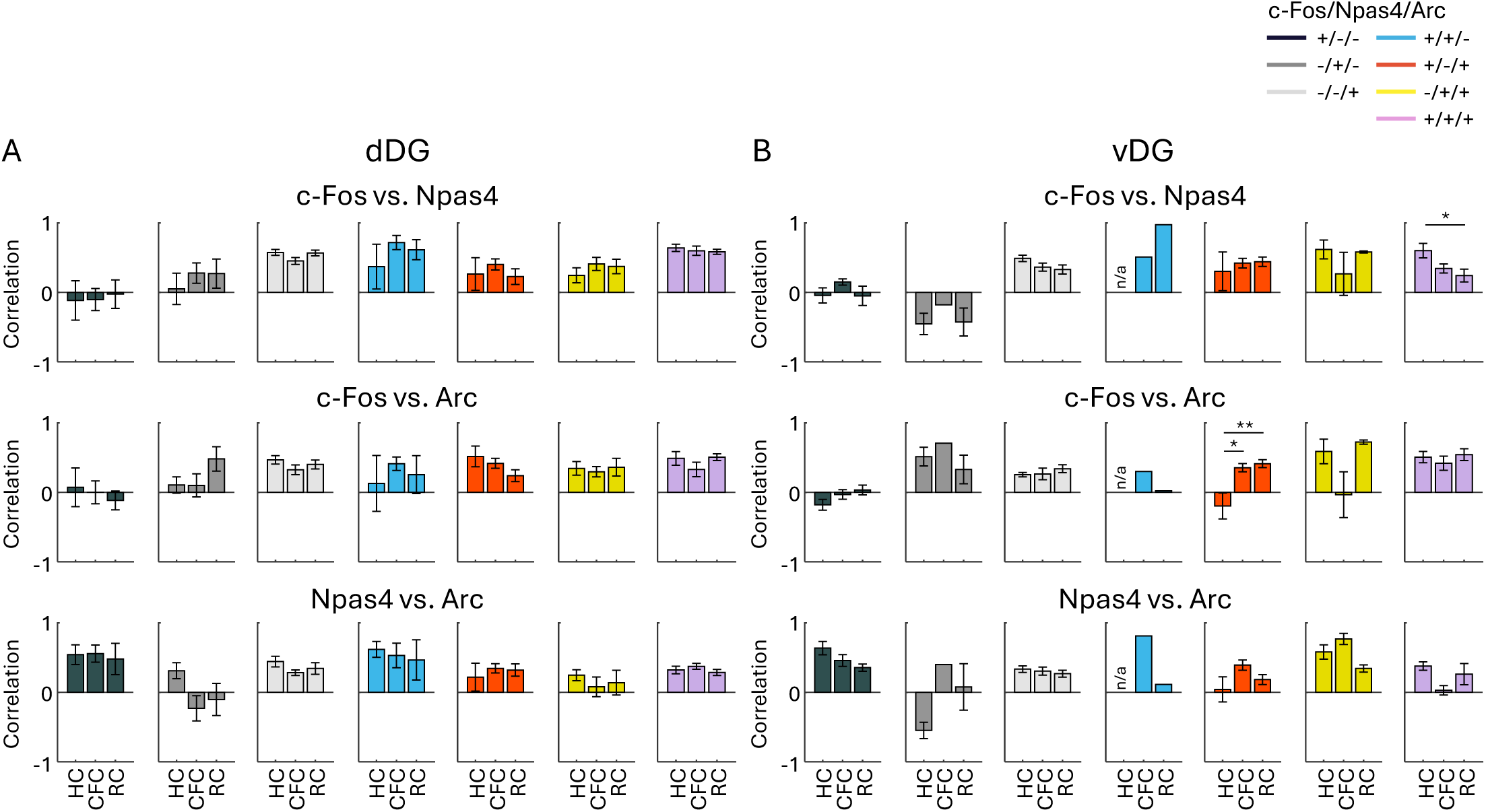
Correlation of IEG Intensities in each cell group in DG Average correlation of IEG intensities of c-Fos, Npas4, and Arc in individual cells in HC, CFC, and RC, in the dDG (A) and vDG (B).

**Supp. Figure S20:**
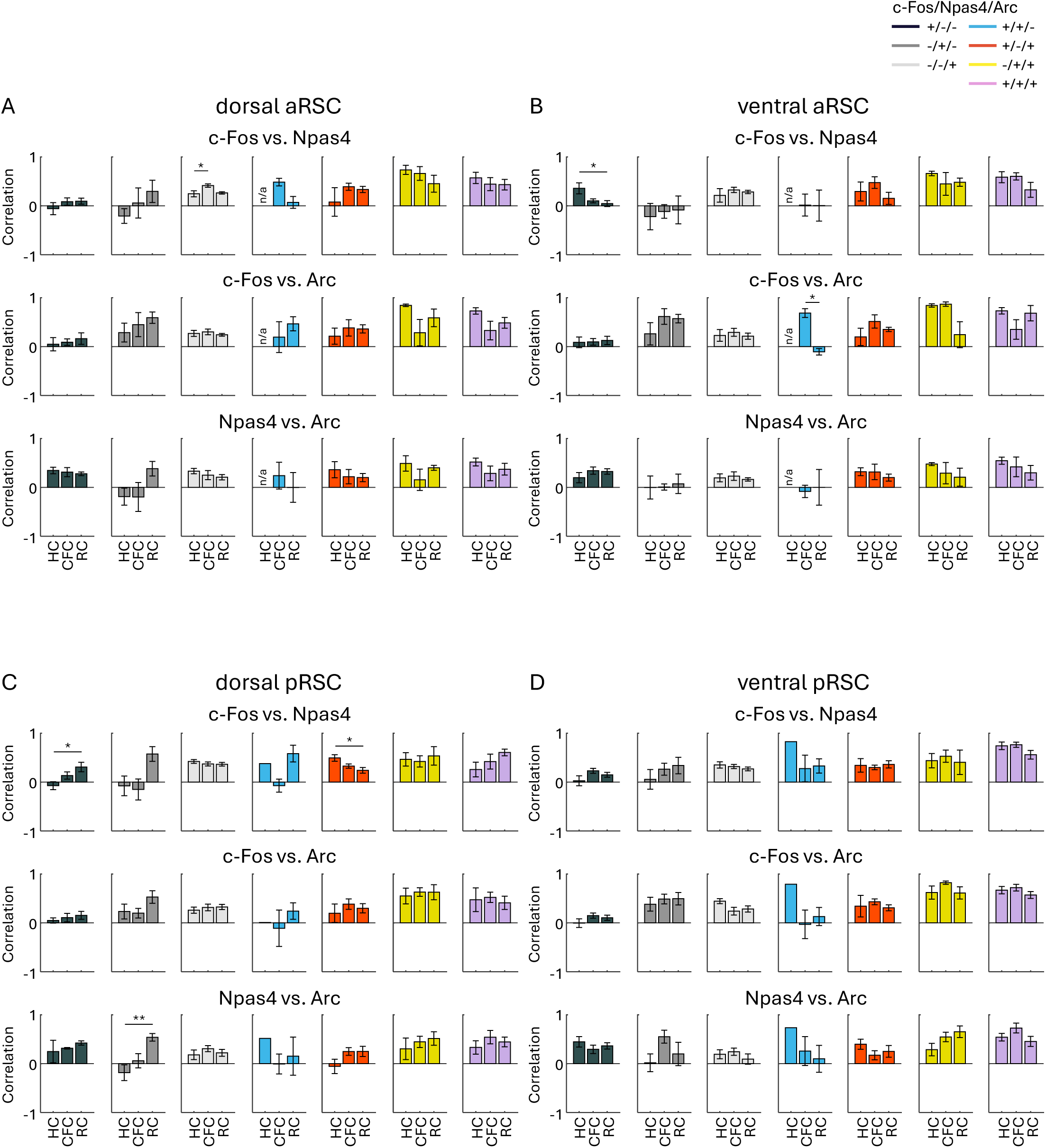
Correlation of IEG Intensities in each cell group in RSC Average correlation of IEG intensities of c-Fos, Npas4, and Arc in individual cells in HC, CFC, and RC, in the dorsal aRSC (A), ventral aRSC (B), dorsal pRSC (C), and ventral pRSC (D).

**Supp. Figure Fig. S21:**
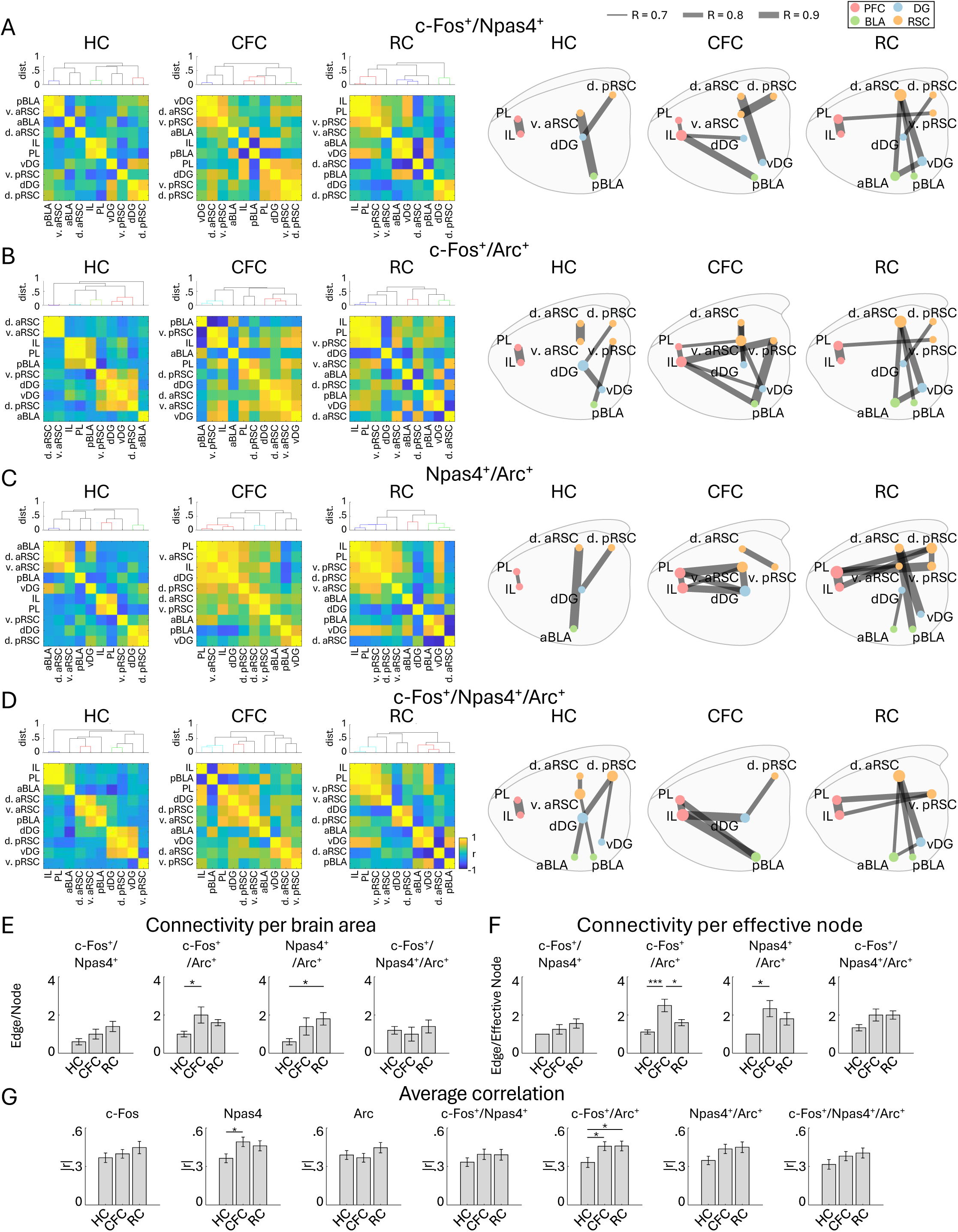
Functional connectivity network of IEG overlapping cells (A–D) Similarly to Figure 8, Inter-regional correlation matrices and connectivity networks based on c-Fos^+^/Npas4^+^ (A), c-Fos^+^/Arc^+^ (B), Npas4^+^/Arc^+^ (C), and c-Fos^+^/Npas4^+^/Arc^+^ cells (D). (E, F) Quantification of network complexity: Average number of edges per node (E) and Average number of edges per effective node (F). (G) Average of absolute correlation values in the correlation matrices for each cell group which are shown in Figure 8A–C and S21A–D.

## Notes

### Competing Interest Statement

The authors have declared no competing interest.

### Summary of Updates

We corrected information of funding supports in Acknowledgements.

